# Searching for Structure: Characterizing the Protein Conformational Landscape with Clustering-based Algorithms

**DOI:** 10.1101/2023.09.13.557631

**Authors:** Amanda C. Macke, Jacob E. Stump, Maria S. Kelly, Jamie Rowley, Vageesha Herath, Sarah Mullen, Ruxandra I. Dima

**Affiliations:** Department of Chemistry, University of Cincinnati, Cincinnati, OH 45221; Department of Chemistry, Emory University, Atlanta, GA 30322; Department of Chemistry, The College of Wooster, Wooster, OH 44691; Department of Chemistry, Virginia Tech, Blacksburg, VA 24061

## Abstract

The identification and characterization of the main conformations from a protein population is a challenging, inherently high-dimensional problem. We introduce the Secondary sTructural Ensembles with machine LeArning (StELa) double clustering method, which clusters protein structures based on the underlying Ramachandran plot. Our approach takes advantage of the relationship between the phi and psi dihedral angles in a protein backbone and the secondary structure of the protein. The classification of states as vectors composed of the clusters’ indices arising naturally from the Ramachandran plot, followed by the hierarchical clustering of the vectors, enables the identification of the minima from the corresponding free energy landscape (FEL) by lifting the high structure degeneracy found with existing approaches such as the RMSD-based clustering GROMOS. We compare the performance of StELa with not only GROMOS but also with CATS, the combinatorial averaged transient structure clustering method based on distributions of the phi and psi dihedral angle coordinates. Using ensembles of conformations from molecular dynamics (MD) simulations of either intrinsically disordered proteins (IDPs) of various lengths (tau protein fragments) or from local structures from a globular protein, we show that StELa is the only clustering method that identifies nearly all the minima from the corresponding FELs. In contrast, GROMOS yields a large number of clusters that cover the entire FEL and CATS, even with an additional clustering step, is unable to sample well the FEL for long IDPs and for fragments from globular proteins as it misses important minima.

**TOC Graphic:** 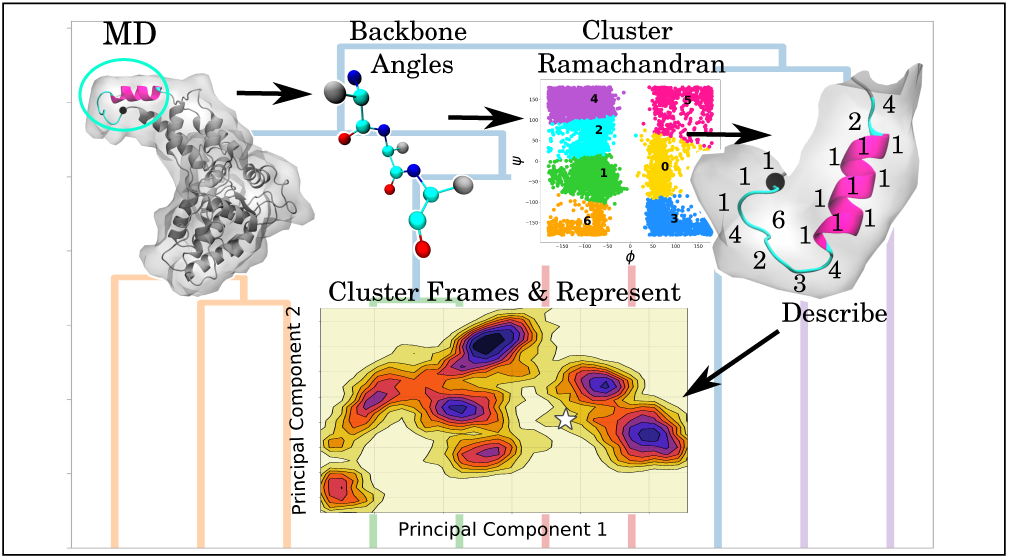

## Introduction

In many enzymes conformational changes can occur due to the binding of ligands, other proteins, or protein mutations. These are fundamental processes with important functional implications.^1^ Such conformational changes sometimes correspond to a global rearrangement of the protein. This is observed in the quaternary structure of the microtubule (MT) tubulin dimers where the straight GTP-bound tubulin becomes curved when the hydrolyzable GTP is chemically converted into GDP.^2–4^ Other times, they correspond to local changes at the secondary level or to a transition from ordered to disordered (or vice versa) as seen in MT-severing enzymes when the substrate binding loops become ordered in the presence of the MT substrate.^1,5,6^ Great effort has been expended to predict the steps that lead to these conformational changes and the corresponding states. Experimental methods, such as X-ray crystallography and cryogenic electron microscopy (cryo-EM), are useful tools for identifying conformational transitions by solving structures in the presence or absence of cofactors of interest.^7^ Computational tools, such as the bioinformatics-based PSIPRED and the AI-based AlphaFold, are employed to predict protein structures from protein sequences using machine learning.^8–12^ However, all of these methods struggle to characterize regions or entire proteins that are “fuzzy”, or that are heterogeneous due to a lack of order and to capture their structural transitions over time.

All-atomistic computational tools, such as MD simulations, allow for a more detailed exploration of the conformational landscape. The challenge then becomes the extraction of the most dominant sampled states from a large and complex data set, particularly when characterizing how environments or allosteric modulators affect the conformational space. The aforementioned proteins that lack well-defined structure are known as intrinsically disordered proteins (IDPs). Despite the usage of the term “disorder,” IDPs do not sample their conformational space in a completely random manner;^13^ an IDP favors certain conformations over others, albeit to a much less extreme degree than structured globular proteins.^14^ IDPs sample a large number of diverse conformations while carrying out their functions in the cell, in apparent contrast to the traditional “structure-function” paradigm.^15–17^ A well-known example of an IDP is the MT-stabilizing protein found primarily in neurons called tau.^18^ Tau is made up of an N-terminal projection domain, a proline-rich domain, a combination of repeat domains (R1-R4 and a pseudo-repeat R’), and a C-terminal domain that are spliced together in various isoforms. The misregulation of tau is associated with neurodegenerative diseases such as Alzheimer’s disease where tau is found to aggregate as a result of a change to its conformational space.^18–22^ Describing these changes is an essential piece for understanding neurodegenerative diseases and therefore MD is commonly employed to probe the underlying conformational space of IDPs.^17,20,23^

IDPs are not the only types of proteins characterized by a large or changing conformational space, which could be challenging to explore. Regions of proteins that experience conformational changes due to the allosteric influence of a modulator are also difficult to characterize. These types of transitions have been captured with the above mentioned experimental methods, as well as with computational tools. In our previous work, using all-atomistic MD simulations to study lower order oligomers of the MT-severing enzyme katanin, we identified a ligand-dependent conformational transition in a region of the protein from a Loop-Helix to a Helix-Helix structure.^24^ As the Helix-Helix structure was also proposed based on the X-ray structure of the monomeric form of katanin,^25^ our finding of the conformational switch in simulations was essential for understanding the allosteric contribution of the binding cofactors in katanin, and for assessing the stability of the lower order oligomers. Additional transient and flexible structures were identified in dimers and trimers in the presence and absence of the binding ligands. MD simulations were also used to identify a nucleotide-dependent conformational transition in secondary structure from a loop in the G-actin filament to a helix in the F-actin filament.^26–28^ Hemoglobin is a classic example of extensive ligand-induced allosteric control over a quaternary protein structure.^29^ A single point mutation (E6V) in the *β* chains of hemoglobin leads to sickle cell anemia.^30,31^ This mutation causes the aggregation of hemoglobin fibrils in the absence of *O*_2_ (T) that leads to the deformation of red blood cells that disintegrate when *O*_2_ is present (R).^31,32^ Having a means of characterizing the conformational landscape is crucial for identifying states for these more heterogeneous proteins/regions, which are essential for understanding enzymatic mechanisms, or the effects of allosteric modulators and of disease-related mutations.

Unsupervised machine learning methods such as clustering algorithms have been used to extract significantly populated states from the sampled space by grouping together structures identified as similar according to a given criterion. The standard for clustering MD conformations data is the root-mean-squared-deviation (RMSD) based algorithm that determines similarity using an RMSD cutoff. Structures within this cutoff are grouped together, whereas a neighbor outside of the cutoff is assigned to a distinct group.^14^ Cheung et al suggested that the RMSD-based clustering might not be descriptive enough for identifying local structures with similar patterns, especially in IDPs. They introduced a new algorithm, CATS (Combinatorial Averaged Transient Structure), which first characterizes structures based on an internal collective variable before determining similarity.^33^ In our previous study, described above, of oligomeric species from MT severing enzymes, we used our in-house clustering algorithm, StELa (Secondary sTructural Ensembles with machine LeArning), to successfully probe for a secondary structural transition in a local region referred to as the helical bundle domain tip (”HBD tip”).^24^ Here, we put our algorithm to the test to map out its performance on a number of protein systems. Namely, we performed clustering of the states from the sampled conformational space of R2/tau fragments, R4-R’/tau fragments, and the HBD tip fragments described above, all taken from MD simulations, using each of the three algorithms: RMSD-based, CATS, and StELa.^14,20,24,33^ We evaluated the differences in the free energy landscapes (FELs) between short and long fragments of IDPs (R2/Tau and R4-R’/Tau), as well as between IDPs and the transient, but ordered, region from a globular protein in a tertiary and quaternary assembly (HBD tip of each of the katanin protomers). Mapping out the clustered states on the FEL in the principal components (PCs) space allowed us to evaluate the ability of each method to sample the conformational space. We conclude with a comparison of the advantages and the pitfalls of each of the methods.

## Methods

### Molecular Dynamics Simulations

To address how the 3 clustering methods discussed in the **Introduction** handle different systems, we used MD simulations for a short fragment of an IDP (R2/Tau),^20,33^ a longer fragment of an IDP (R4-R’/Tau), a short fragment from a tertiary globular protein structure (katanin), and the same fragment from its quaternary form (ABC/katanin).^24^ The MD simulations for the R2 fragment of Tau (14 units), computed with the GROMACS software package, detailed in Table S1, were from an earlier paper. Here, the authors tested how different solvents induce the formation of aggregation-prone states of R2/Tau: Urea to mimic a denatured state, Water for a more standard state, and TMAO for a more aggregation-prone state.^20,34–36^ The models used for TMAO (2M) and Urea (5M) were previously developed by Weerashinghe and Smith, and Larini and Shea.^37,38^ For their simulations, the authors used the all-atom optimized potentials for liquid simulation (OPLS) force field with three-site transferable intermolecular potential rigid water.^20,39,40^ The structural changes observed in these simulations, resulting from an altered conformational landscape, were characterized by Cheung et. al using two different clustering methods: the RMSD based algorithm and their in-house developed algorithm, CATS.^33^

We carried out all-atom MD simulations for R4-R’/Tau (121 residues) using GROMACS 2022 and the AMBER99SB-ILDN force field. ^34–36,41^ We created the structure used in the simulations by applying the Modeller plug-in from the ChimeraX visualization software to the PDB 7PQC configuration.^42–44^ We then centered this structure in a dodecahedral box with periodic boundary conditions (PBC), solvated with TIP3P water molecules, and neutralized with NaCl.^45^ We performed energy minimization for 50,000 steps using the steepest descent algorithm and the Verlet cutoff scheme.^46^ We did NVT equilibration for 500ps, followed by NPT equilibration for an additional 500ps. Both equilibrations employed the leapfrog integrator, Verlet cutoff scheme, and the velocity-rescaling thermostat. The NPT equilibration used also the Parrinello-Rahman pressure coupling.^47,48^ We ran four production trajectories, each 200ns long, using the same temperature and pressure coupling methods as in the NPT equilibration. Coulombic interactions were treated using the Particle Mesh Ewald (PME) algorithm, while other non-bonded interactions were computed using the Verlet cutoff scheme.^46,49^ The RMSD over time of these new trajectories, shown in Figure S1, was used to monitor the equilibration of the system.

The MD simulations, carried out with the GROMACS software package using the GROMOS96 54a7 force field and SPC solvent, used for the katanin analysis, detailed in Table S2, were taken from our previous studies where we investigated the conformational behavior of lower order oligomers, proposed to form upon disassociation of the fully functional hexamer at the start of the MT severing process.^24,34,36,50,51^ It is important to note that the spiral conformation corresponds to the pre-hydrolysis assembly of katanin, characterized by a gap between the terminal protomers (A and F). In turn, the ring conformation is a post hydrolysis state where the nucleotide is missing from protomer A resulting in the closing of the gap between the end protomers.^5,6^ In our study we probed monomers from the spiral conformation, as well as dimers (AB, BC) and trimers (ABC) from the spiral and ring conformations of katanin, as indicated in Figure S2. To understand the effects of binding the ligands, the monomer was simulated with both the ATP and polyglutamate MT minimal substrate (COMPLEX), as found in the solved cryo-EM structure,^6^ with ATP only (NUC), with the minimal substrate (SUB), and in the absence of both ligands (APO). The trimers were simulated in the COMPLEX and APO states. We observed that a region in the helical bundle domain, which we dubbed the “HBD tip” (see Figure S2), consisting of amino acids ASP417 to LEU437 (21 peptide bonds), showed significant structural fluctuations in several of the setups. Our hot spots analysis determined that only 3 regions from katanin experience allosteric changes upon the binding of the ATP, the MT substrate, and the formation of the various interprotomer interfaces. The HBD tip was one of these regions.^24^ We developed our in-house algorithm, StELa, to characterize such structural changes from the extracted HBD tip region conformations of each protomer in a simulation. Clustering with StELa showed that this region undergoes a structural change in the monomer upon the binding of the ATP and of the minimal MT substrate. In the ring ABC-trimer, we found a similar structural change associated with the binding of cofactors as seen in the monomer. However, this time the concave interface made between the *i*th and the (*i* + 1)th protomers and/or the convex interface formed between the *i*th and the (*i −* 1)th protomers, described in Figure S2, also caused interesting changes to the conformational space of the HBD tip in the i-th protomer (B).^5,6^ In this oligomer, protomer A has only the concave interface, formed between both its Nucleotide Binding Domain (NBD) and Helical Bundle Domain (HBD) regions and the NBD of protomer B. Protomer C has only the convex interface, formed between its NBD region and the NBD of protomer B. Our analysis of the PC motions showed that protomer A is highly flexible, due to its absence of the nucleotide, so much so that its NBD detached from the NBD of protomer B. We determined that, while protomers B and C would likely remain bound to each other, protomer A would dissociate from protomer B due to the motions leading to the detachment of its HBD from the NBD of protomer B. This indicated that the NBD-HBD contacts are more important for the stability of the interprotomer interface and the overall stability of the machine than the NBD-NBD interfaces. Importantly, the NBD-HBD contacts are formed with the long HBD helix and the loop/helix of the HBD tip.^24^

### Current Methods for Clustering Protein Fragments

The GROMOS algorithm implemented with GROMACS has long been considered the gold standard for evaluating the types of structures explored in MD trajectories.^14,34,36^ This algorithm uses the RMSD to evaluate how similar a data point, or structure, is to its neighbors, within a chosen cutoff (0.14 nm).^14,33^ This means that the clustering is based on the Cartesian coordinate space of the structure. The cluster centers, which are used as cluster representatives, are the structures with the largest number of neighbors. Cheung et al used this method as a control for testing the performance of their in-house algorithm, CATS (<u>C</u>ombinatorial <u>A</u>veraged <u>T</u>ransient <u>S</u>tructure).^33^ The CATS algorithm (https://github.com/Cheung-group/CATS) converts the structure found in each frame of a trajectory to a vector, prior to determining similarity using clustering. This time, the authors chose the phi and psi dihedral backbone angles as descriptive collective variables (CVs), instead of the Cartesian space-based CVs used by GROMOS.^33,52^ This was significant, as the dihedral angles provide a detailed internal descriptor of the secondary structure of proteins and are known to capture conformational changes.^53^ The user provides the phi and psi backbone angles as the input, which we extracted using the **rama** function in GROMACS. Distributions of each residue’s phi and psi angles are fitted to Gaussians in order to identify one to three main peaks. Next, CATS creates representative vectors for each trajectory frame consisting of a series of integer labels corresponding to the Gaussian curve each angle was found in, resulting in two labels per residue (Figure S3). CATS then groups together identical representative vectors, which could result in a large number of small clusters, as seen with the RMSD-based algorithm; however, this makes it difficult to analyze/identify the types of important structures sampled by the simulations. To address this issue, we added a centroid-based clustering step at the end of our implementation of CATS, which we refer to as “CATS+”, to group similar clusters. This also makes for a more fair comparison between the CATS+ and StELa results. Our version of CATS was written in Python, while the original CATS algorithm used a combination of MATLAB, C++, and tcl.

### Clustering Protein Fragments with StELa

Inspired by the use of phi and psi dihedral backbone angles as a collective variable for describing the structure before determining similarity, we have developed our own in-house algorithm, StELa (<u>S</u>econdary s<u>T</u>ructural <u>E</u>nsembles with machine <u>L</u>e<u>A</u>rning). The first version of this algorithm was described in our previous work where we used StELa to characterize the conformations of the HBD tip in katanin.^24^ StELa is a double clustering algorithm, written in Python, which uses libraries such as SciPy and sklearn.^54,55^ Similar to CATS, StELa first characterizes the input structures per sampled frame using the calculated phi/psi backbone torsion angles extracted from the MD trajectories. This set of angles is the ideal descriptive reaction coordinate for characterizing types of protein structures because it defines the geometry observed in specific secondary structure motifs.^33,56,57^ These backbone angles are plotted together on the Ramachandran plot, a classic tool for describing secondary structure. It has been well established that residues in secondary structures populate specific regions of this plot, in particular alpha helices and beta strands.^52,58^ We take advantage of this by applying the first round of clustering directly to the phi/psi dimensions with the centroid based k-means algorithm. In doing so, we determine the number of clusters based on whether or not they split these well-defined regions associated with alpha helices and beta sheets as described in Figure S4. We then convert each structure from a given sampling of frames from MD into a representative vector composed of the cluster labels describing each region of the Ramachandran plot. In doing so, we describe the local structure of a residue in the protein fragment with a single integer. In the previously reported algorithm, we then performed an automated biochemical correction of the vector geometries to ensure that an alpha helix cannot be shorter than 4 consecutive residues with at least 2 consecutive residues between separated helices.^56^ In addition, we now apply a similar check to the beta sheet region of the plot, such that beta strands cannot be shorter than 3 consecutive residues.^57^ To this corrected set of representative vectors, we then apply complete linkage agglomerative hierarchical clustering with Euclidean distance metric as our second clustering step. In complete linkage agglomerative clustering, structures are sequentially grouped into larger clusters based on the maximum distance between elements. The number of clusters was chosen using statistical-based measurements called the silhouette score, calculated with Equation S1, and the Calinski-Harabasz index, calculated with Equation S2.^54,59–61^ The representative structure is chosen by calculating the most probable vector per position for a given cluster and then identifying a matching frame.

### Free Energy Landscapes

In order to test the performance of each method, we extracted the representative structure for each cluster and represented it on the Free Energy Landscape (FEL).^21^ The FEL was previously expressed in the *R_G_*/*R_EE_* space, as seen in Figure S5a; however, it was determined that the minima in the *R_G_*/*R_EE_* space lumps unique structures into the same minima. ^33^ Here, we chose to plot the FEL in the principal component 1 (PC1) and 2 (PC2) space (evaluated with GROMACS **anaeig**), following our previous study.^50^ As shown for example in Figure S5b, this choice of FEL coordinates identifies an additional minima that is not observed in the *R_G_*/*R_EE_* space with a clear barrier between them.^34,36,62,63^ The FELs for the described systems are shown in Figures S6-S8 for comparison. The variance covered by the discussed PCs for each system is in Table S3. The included variance for R2/Tau covered around 30%, for R4-R’/Tau covered over 40%, and for the HBD tip of katanin it covered over 50%. These FELs provide a particularly useful tool for evaluating the differences between the three methods. By plotting the cluster centers or representatives on the corresponding FEL, we can compare the conformational spaces captured by each method. It is important to note that we do not apply clustering to the FEL as done in other studies.^31,64^

## Results

### StELa selects an optimal number of clusters by using statistical checkpoints in a second clustering step to determine the number of unique structures

One of the main challenges encountered when using clustering algorithms, which stems from their unsupervised learning origin, is the decision on the number of selected clusters. For example, in the case of secondary structures, this means how many unique structures are found in the dataset. In our StELa algorithm this challenge appears in the second step, when using the complete linkage hierarchical clustering. To address this challenge, we relied on statistical-based measures such as the silhouette score and the Calinski-Harabasz index which yield a score versus the number of clusters.^54,59–61^ A maximum (usually local) in either of these two scores signals the number that results in the best separation of the clusters, thus indicating the optimal number of clusters. More often than not we found the two scoring methods to be in agreement in their selection of the number of clusters, as seen in Figure S9. An additional checkpoint consists of the analysis of the corresponding dendrogram, an example of which is shown in Figure S10, to determine if the breakdown of the selected number of clusters agrees with the organization of the dendrogram.^54^ The final checkpoint consists of the analysis of each of the resulting clusters to ensure that the selected number of clusters adequately separates unique structures. The number of clusters identified by each algorithm for the various systems probed is reported in Tables S4-S7. Across all of the katanin protomers included in this study, we found the number of clusters to be around 13 for the HBD tip. In turn, for the R2/Tau we found around 20 clusters, and for R4-R’/Tau we needed 23 clusters to describe the respective conformational spaces. Our results show that, in general, the more disordered the fragment, the higher the number of clusters that gets selected by StELa. Importantly, for all the systems, the number of clusters is small enough to make it manageable for future analysis. In turn, the RMSD-based algorithm for katanin identified around 30 clusters for each of the monomers, around 20 clusters for protomers A and B, regardless of the presence of the cofactors, and around 75 clusters for protomer C. For R2/Tau, in each of the solvent conditions, the RMSD-based algorithm identified over 1500 clusters, which is untenable for further analysis. Finally, the CATS+ algorithm selected less than 15 clusters across all of the systems probed in our study. Next, we discuss in detail the results for each protein system probed with the 3 clustering algorithms.

### Selection of Representative States from Small Intrinsically Disordered Protein Fragments: R2/Tau

Previous work on the R2 fragment of tau focused on studying the effects of different solvents on the conformational space of this peptide, which can influence its aggregation propensity.^20,33^ The FELs in the PC space, shown in Figure S6, provide insight into how the different solvents affect the conformational landscapes. We found that all of the landscapes are characterized by one main basin with multiple minima separated by low energy barriers. In water (a) 3 main minima were identified, separated by comparatively higher barriers than those found for the TMAO (b) or urea (c) solvents. R2/Tau in TMAO, the solvent used to induce aggregation, sampled three distinct minima. In contrast, R2/Tau in urea, the solvent used to prevent the formation of secondary structure, was largely confined to a single minima.

The clusters found with each clustering method for the R2/Tau fragment in water are shown in Figure 1. We located five potential minima of interest from the FEL, indicated in panel (a), with the corresponding structures shown in the (i-v) snapshots. We identified two deeper minima, one in the upper region (i), which corresponds to a bent loop, and one towards the middle region (v), which corresponds to a loop with some helical characteristics.

**Figure 1:**
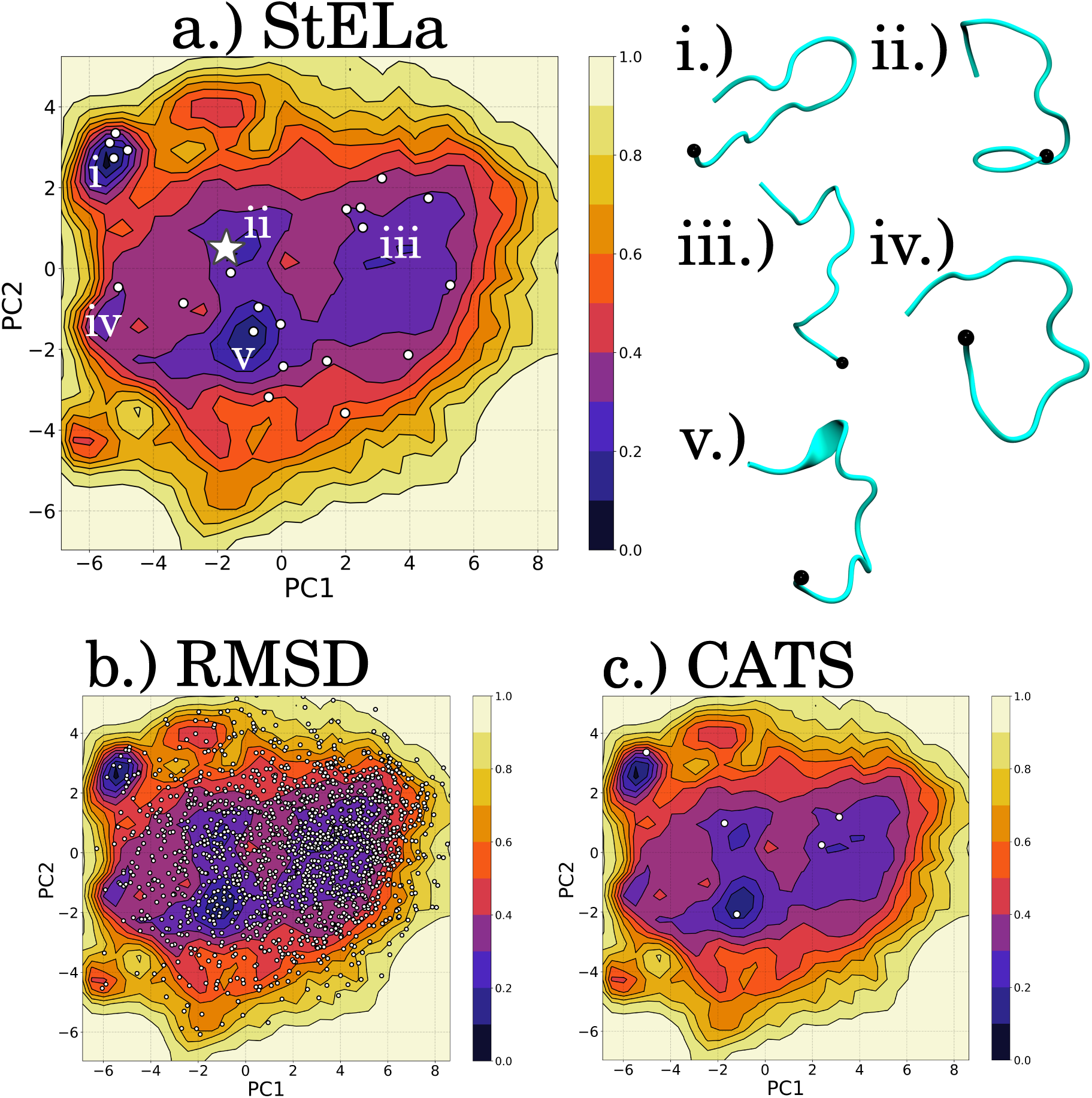
The representative structures for the identified clusters of the R2 fragment of Tau in water plotted on the FEL in PC1/PC2 space for each method: (a) StELa, (b) RMSD and (c) CATS+. The white star in (a) indicates the structure at the beginning of the simulation. Representative structures (i-v) were extracted to represent the minima indicated in (a). The N-terminal end of the structure is indicated with a black bead.

The RMSD-based algorithm, depicted in panel (b), struggled to find any similarity between the probed loop structures, resulting in a large number of clusters.^33^ The CATS+ algorithm identified all but one minima; however, the resulting populations, shown in Table 1, used to represent the two deeper minima are *<* 1% for (i) and (v), which is unexpected considering their low energy. The majority of the population is located in the clusters from region (iii), with the remaining population contributing primarily to region (ii). The StELa clusters identified each of the five regions and with populations closer to the expected range, such that the majority of the structures populated regions (i) and (ii). The remaining population was located in clusters representing structures from higher energy states, rather than from any of the identified minima. The above trends found in the RMSD-based and CATS+ algorithms persisted across the different solvent systems, as shown in Figure S11 for TMAO and in Figure S12 for urea. Interestingly, the FEL from the TMAO setup showed that region (i) corresponds to a folded *β*-sheet. CATS+ did identify a cluster near this area that also showed the folded *β*-sheet conformation; however, the population was 0.3%, whereas StELa found it to be 15%. The most striking difference between the clustering for R2/Tau according to the 3 methods was in the populations of the clusters, most notably of the largest cluster, as detailed in Table S4. The population of the largest cluster for the RMSD-based algorithm was 4%, for CATS+ was 85%, and for StELa was 28%. This cluster for the RMSD-based and StELa algorithms is located in region (ii), while for CATS+ it is found in region (iii). These results illustrate the difficulty in grouping together structures from the MD datasets.

**Table 1:** The collective populations for the clusters found in each minima from Figure 1.

| **FEL Region** | **CATS+** | **StELa** |
| --- | --- | --- |
| **i** | 0.2% | 20% |
| **ii** | 5% | 28% |
| **iii** | 95% | 9% |
| **iv** | - | 0.8% |
| **v** | 0.3% | 12% |

### Selection of Representative States from Large Intrinsically Disordered Protein Fragments: R4-R’/Tau

We also tested the performance of the clustering algorithms when applied to a longer IDP. For this, we selected a 121-residue long portion of tau, consisting of the fourth repeat in the MT binding domain and the adjacent pseudo-repeat located towards the N-terminal. The FEL calculated for this system (Figure 2) shows four minima (i-iv), with minima ii and iii located in a shared basin and separated by a relatively low energy barrier. Minima (i) and (iv) are separated from this basin and each other by larger barriers. This is noticeable when compared to the shorter Tau/R2 system, whose FEL exhibited only low energy barriers between the minima. Figure 2 depicts the cluster centers/representatives for each of the three algorithms plotted on the FEL. This clearly shows that CATS+ finds only a single cluster with a representative frame in minimum (iii). The RMSD-based algorithm finds thousands of clusters, but the centers for these clusters reside only in the large basin containing minima (ii) and (iii). In contrast, StELa is the only algorithm able to find clusters that cover all four minima in the FEL.

**Figure 2:**
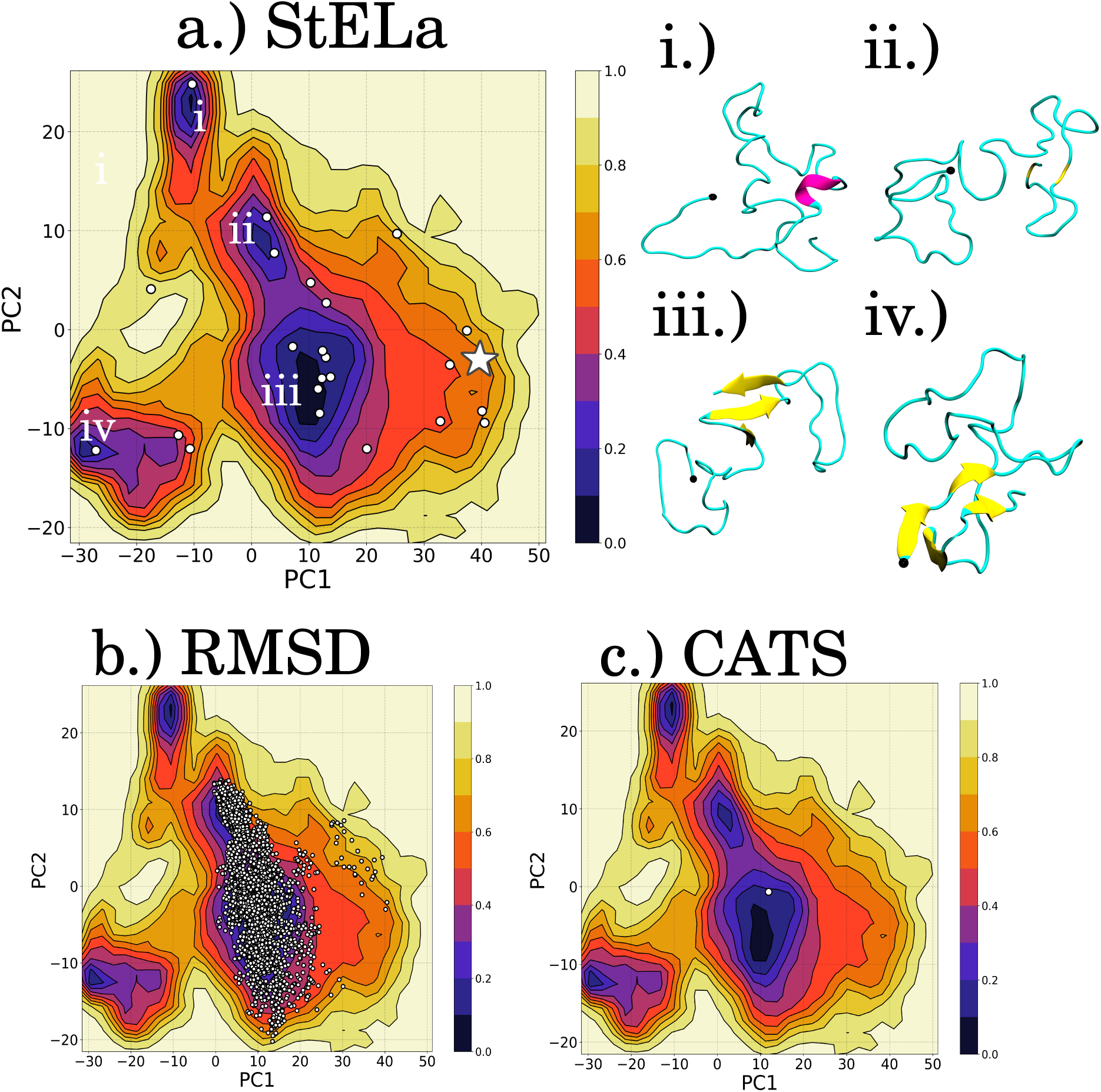
The representative structures for the identified clusters for the R4-R’ fragment of Tau in water plotted on the FEL in PC1/PC2 space for each method: (a) StELa, (b) RMSD and (c) CATS+. The white star in (a) indicates the starting structure at the beginning of the simulation. Representative structures (i-iv) were extracted to represent the minima from (a). The N-terminal end of the structure is indicated with a black bead.

### States from a Tertiary Protein Fragment: the HBD tip of the Katanin Monomer

In a previous study, we determined the effects of binding the associated cofactors, ATP and the MT minimal substrate, on the structural stability of lower order oligomers of katanin.^24^ The HBD tip fragment, taken from the tertiary structure of the katanin monomer as described in the Methods, has well-defined FELs in the various setups (Figure S7). The fragment from the monomer in the NUC setup (b) resulted in relatively more shallow energy barriers in comparison to the other setups, but still high in comparison to the R2/Tau FEL. The FEL for the COMPLEX (a) and SUB (c) setups resulted in minima separated by high and wide energy barriers along the PC1. We noticed that the FELs for these katanin monomers have a similar number of well defined regions, with the change in energy barriers reflecting their dependence on the presence of the cofactors.

We found six minima of interest for the COMPLEX setup of the katanin monomer, as indicated in Figure 3a, with the identified representative structures in (i-vi). The deepest minima are characterized by (i) and (iii) whose structures are differentiated by the length of the helix and the orientation of the loop region. A more shallow region (ii) corresponds to the C-terminal helix being similarly long as in region (iii), but the orientation of the N-terminal loop being more similar to that found in region (i). This is intriguing because the clustering identifies a region of the FEL between the two minima which seems to correspond to a potential intermediate state between (i) and (iii). There is a large and wide barrier described by PC1 to another group of minima in the landscape where structures (iv - vi) are found. The structure in region (iv) corresponds to a state where the N-terminal loop transitioned into a 3-10 helix according to the STRIDE assignment with VMD; however, as previously mentioned, StELa defines helices as 4 helical angles in a row.^65^ The structures that represent (v) and (vi) correspond to loop regions and a helix of varying length. For this well-defined tertiary structure of the fragment from the katanin monomer, the RMSD-based algorithm (b) finds a smaller number of clusters in comparison to those found for Tau, as seen in Table S6, but those clusters only identify the minima corresponding to structure (iii). CATS+ only identifies 3 clusters, all of which represent the same minima that corresponds to structure (i). For the other setups of the katanin monomer, shown in Figures S13-S15, similar sets of representative structures were identified corresponding to the various minima. The RMSD-based algorithm sampled two of the minima found for the NUC setup, one for the SUB setup, and four for the APO setup. CATS+ identified three minima for the NUC setup, two for the SUB setup, and two for the APO setup. The largest cluster for the RMSD-based algorithm, corresponding to region (iii), was 38%, for CATS+ corresponding to region (i) was 56%, and for StELa corresponding to region (iii) was 30%. StELa only failed to characterize one minima in the APO setup out of the systems for the katanin monomer.

**Figure 3:**
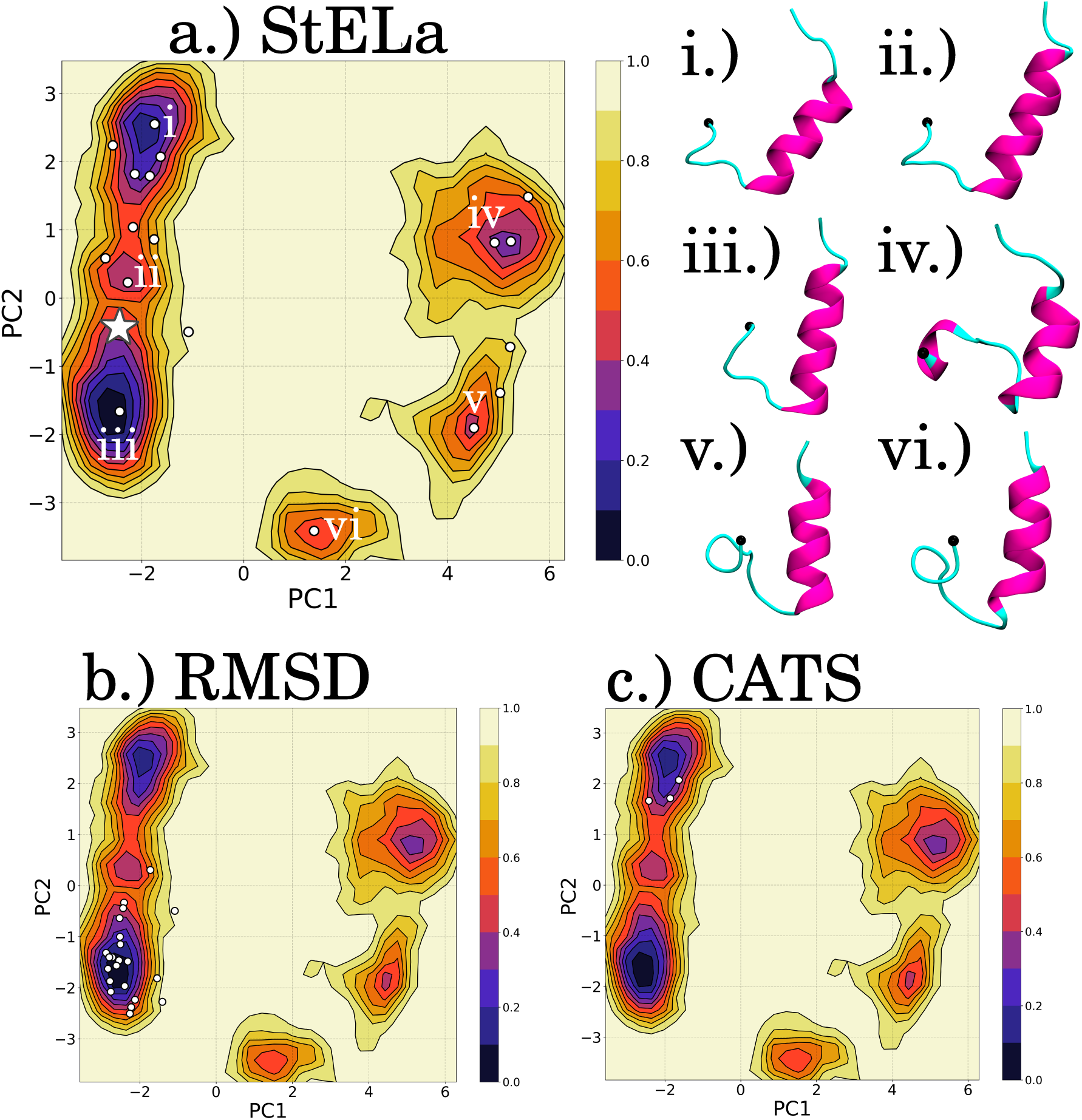
The representative structures for the identified clusters for the HBD tip of the Monomer COMPLEX set up plotted on the FEL in PC1/PC2 space for each method: (a) StELa, (b) RMSD and (c) CATS+. The white star in (a) indicates where the starting structure is at the beginning of the simulation. Representative structures (i-vi) were extracted to represent the minima indicated in (a). The N-terminal end of the structure is indicated with a black bead.

### Selection of Repsentative States from a Protein Fragment in Quaternary Assemblies: the HBD tip of each Protomer of the Katanin Ring Trimer

In our previous study, we were also interested in characterizing the effects of the interprotomer interface formation (i-1,i and i+1,i) on the conformational flexibility and allostery of severing enzyme protomers in dimers and trimers.^24^ We used the average RMSD and the PC motions to compare the stability of each of these lower order oligomers. Our analysis showed that the ring ABC trimer was the least stable quaternary configuration, characterized by the dissociation of protomer A due to the increase in disorder of its NBD due to the lack of the nucleotide.^5,6^ This movement resulted in the the bending and conformational changes in the HBD tip of protomer B, which in turn allowed it to preserve its contacts with protomer C. The lack of the concave interface in C led to a mobile HBD and more flexible structures in its HBD tip. The characterization of this region in each of the protomers was particularly important for understanding the inner working of the katanin oligomers: the persistence of contacts between the HBD tip fragment of protomer i and the NBD of protomer i+1 is essential for the stability of any quaternary assembly. It thus comes as no surprise that the FELs for each protomer of the ring ABC-trimer in the COMPLEX and APO setups are distinct depending on the bound cofactors and the presence of different types of interfaces (Figure S8). Moreover, these profiles are also distinct from the corresponding FELs of the katanin monomer (Figures S7), which supports the idea that the HBD tip experiences conformational changes due to the presence of the specific interprotomer interfaces in the quaternary ensembles.

The FEL for protomer B in the APO setup, seen in Figure 4, is particularly interesting due to the unique structures corresponding to a bent helix in the HBD tip. In this FEL, region (iv) contains a structure resembling the starting structure and region (i) corresponds to the formation of a helix in the N-terminal end. Regions (ii) and (iii) are separated by low energy barriers between regions (i) and (iv). The structure associated with region (ii) has a single turn helix in the N-terminal region and a helix in the C-terminal end. The structure associated with region (iii) has a loop in the N-terminal end and a dramatically bent helix in the C-terminal end. As pointed out above based on the putative intermediate state in the COMPLEX setup of the monomer, these regions likely correspond to two intermediate states between the Loop-Helix structure and the Helix-Helix structure. Region (v) is broad, but relatively shallow compared to the other minima, and it is separated by a high energy barrier from the other regions. This region corresponds to a flexible structure, with a single turn helix in the middle of the fragment. The RMSD-based algorithm (b) only identified region (iv), while CATS+ (c) identified clusters from (i), (iv) and (v), but did not identify the described intermediate regions (ii) and (iii). In contrast, StELa (a) identified all of the regions. The results obtained using StELa were key in determining that the unique behavior of the conformational space of protomer B is due to the permanence of its contacts with the NBD of protomer C.^24^ Moreover, this illustrates the important role that an accurate representation of the unique types of secondary structures has in gaining an understanding of complex biological machines. In the COMPLEX state, shown in Figure S18, the FEL for protomer B presents five regions of interest. Regions (i) and (ii) are close to each other in space and configurations, being separated by medium energy barriers, corresponding to various N-terminal loop orientations and C-terminal helix lengths. Region (v) represents a similar C-terminal helix, but it is higher in energy due to a tight turn and coiled loop arrangement. Region (iv) corresponds to a bent C-terminal helix, similar to that found in the APO set up. StELa identified clusters corresponding to each of the described regions, while the RMSD-based algorithm only identified two and CATS+ identified three regions. In summary, the analysis of the lower-order oligomers provided useful insight into the influence of the cofactors, and of the interfaces, for the structural and functional behavior of katanin. A full description and comparison of the results for each of the protomers in the ring trimer in the presence and absence of the binding cofactors can be found in the *Supplementary Information*.

**Figure 4:**
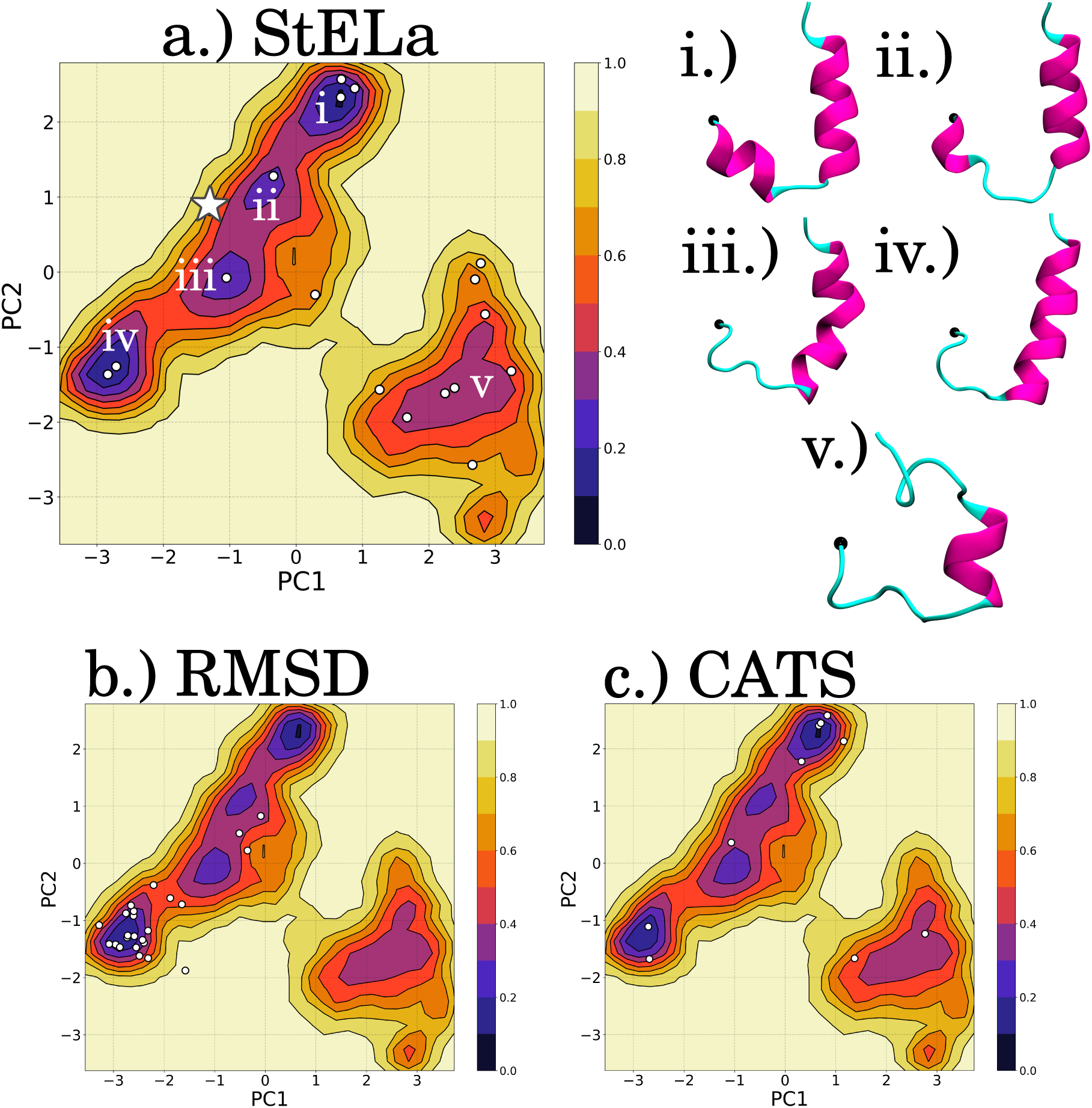
The representative structures for the identified clusters for the HBD tip of Protomer B from the Ring-ABC-APO set up plotted on the FEL in PC1/PC2 space for each method: (a) StELa, (b) RMSD and (c) CATS+. The white star in (a) indicates where the starting structure is at the beginning of the simulation. Representative structures (i-v) were extracted to represent the minima indicated in (a). The N-terminal end of the structure is indicated with a black bead.

## Discussion and Conclusions

### Unsupervised machine learning offers powerful tools for describing and extracting representative protein structures

Computational tools such as MD simulations allow us to access the conformational landscape of proteins in atomistic detail under dynamic equilibrium conditions.^14^ The challenge of identifying the rich structural variety from MD data is due to the sheer size of the system, which makes the identification of the number and the identity of the unique states a very challenging problem. Because usually there is little or no prior knowledge regarding the number of the representative states, unsupervised learning methods such as clustering are the only types of machine learning tools that can be employed to identify subgroups from a dataset.^66^ Characterizing these disordered or flexible regions, often associated with functional regions, allows us to describe the changes to the conformational landscape in different environments or due to ligand binding, which is crucial to understanding enzymatic mechanisms. Accessing these states is a starting point for identifying functionally-relevant conformations, determining a structure appropriate for docking, or for accelerated simulations to further interrogate the conformational space of the protein of choice.^67^ Challenges associated with clustering are the choice of clustering approach for a given dataset, and how to determine the number of clusters/states. Importantly, it is also possible to think of approaches that perform a clustering of the initial set of clusters to further reduce the data complexity and collectively describe the structures from a simulation.^68^

### RMSD based clustering captures large global changes but struggles to determine similarity for IDPs and secondary structures in parts of globular proteins

We found that the standard RMSD-based algorithm favors a large primary cluster that usually gravitates towards the folded native structure, which can be particularly useful for well defined protein structures that generally maintain that same shape, as observed for many tertiary structures of globular proteins.^2,3,14^ This is particularly useful when the goal is to quickly describe global tertiary or quaternary sampled states, as found in previous studies.^14,68^ Unlike globular proteins, IDPs are characterized by broad conformational landscapes and shallow energy barriers. As a result, they tend to populate a rich variety of structures, which makes it difficult for algorithms to appropriately identify similarity between states. The RMSD-based algorithm, which determines similarity based on a RMSD cutoff, has failed to describe short or long IDP fragments, as it produced a large number of small clusters.^33^ This is further demonstrated by the significantly overlapping RMSD distributions we found for R2/tau, shown in Figure S21. The RMSD algorithm has better performance when applied to the HBD tip region from the tertiary and quaternary protein structures and results in similar populations for the largest clusters, as described in Table S6 and S7; however, it only characterized one or two of the identified minima in each system, which were the lowest in energy. While these minima described states considered to be the most populated or native, it failed to describe the full conformational space as it missed or over-represented key regions of interest. The minima that corresponded to the Helix-Helix conformation, that we previously associated with ligand binding, was inappropriately represented by the RMSD-based algorithm (for example: missing in the trimer-A-COMPLEX setup (Figure S16b) and over-represented in the monomer-SUB setup (Figure S14b)). In conclusion, the RMSD-based clustering does not reflect the atomistic detail required for determining similarity in IDPs, as previously reported,^33^ nor for local secondary structures, as seen in the monomers and trimers of katanin.

### Reducing the backbone torsion angles to a single dimension allows for better structural characterization prior to clustering of protein structures

To address the challenge of finding a more useful similarity metric, Cheung et al created the CATS algorithm, based on the phi and psi backbone torsion angles which is a well known and detailed internal descriptor for the secondary structures of proteins.^33^ Prior to determining similarity, the user describes the structure with a chosen number of labels from Gaussian distributions of the phi angle and then the psi angle. The algorithm then defines similarity based on whether or not two vectors are found to be identical, which takes additional time.^33^ The longer the protein, the more time it takes to make these decisions. One of the drawbacks they observed for the clustering of R2/Tau was that, similarly to the RMSD-based algorithm, CATS identifies a large number of small clusters. When an additional clustering step, using k-means, is applied to the resulting clusters, CATS+ identifies no more than 10 similar clusters for the tau datasets. Even more striking is the fact that CATS+ only identified 1 cluster for the longer tau fragment (R4-R’) (Figure 2c). Similarly to the RMSD-based algorithm, we found that the clusters extracted with CATS+ resulted in an inappropriate representation of the regions and, at times, did not identify regions of the landscape at all, as seen in the R2/Tau TMAO set up (Figure S11c). When clustering the katanin datasets, CATS+ provided a relatively reasonable set of clusters compared to the Tau results, but it still struggled with misrepresenting and identifying the minima of interest, as observed in more challenging landscapes from the COMPLEX set up of the monomer (Figure 3c), of protomer B in the APO setup (Figure 4c) and of protomer C in the COMPLEX and APO setups (Figure S19c and Figure S20c). Additionally, one of the reported limitations of CATS was its ability to characterize the regions associated with beta strands and sheets that are found in abundance in the R2/Tau simulations.^33^ The Ramachandran plots for R2/Tau had higher density in the beta region in comparison to the HBD tip of katanin, as seen in Figure S22, signaling the importance of finding a better way to describe beta strands and sheets than the approach employed by CATS. By considering the phi and psi angles together and reducing the two dimensions into one using centroid cluster labels, our StELa approach was able to overcome this shortcoming of CATS in characterizing all the major protein secondary structures.

### StELa identifies unique regions from the free energy landscape for IDPs and secondary structures

Characterizing the protein fragments prior to clustering, as done by Cheung et al, opens up exciting doors for probing local conformational transitions.^33^ Inspired by CATS, in our algorithm we used the Ramachandran plot more holistically to describe proteins and defined helices as at least four helical angles and a beta strand as at least three angles in the beta sheet region. We then determined the number of states for each setup using statistical scoring methods and applied hierarchical clustering directly to these enhanced descriptive vectors. In our algorithm, we employed the maximum Euclidean distance between vectors to signal similarity. The KMeans and hierarchical clustering functions from sklearn and scipy are quite efficient at handling the dataset.^54,55^ The resulting clusters from StELa were striking. The clusters for the tau data sets more appropriately represented and identified unique regions from the FEL in each of the solvent environments. This was particularly true with regard to the significantly longer R4-R’ fragment. Similarly, the clusters identified from the katanin datasets were able to sample the entire landscape, only missing a minima of interest in the monomer-APO setup (Figure S15a), and the trimer-C-COMPLEX setup (Figure S19a), although neither the RMSD-based nor CATS+ captured these two minima. In our previous study, we used StELa to characterize the observed changes in the HBD tip for various lower order oligomers of katanin.^24^ The results of this analysis identified a state that resembled the structure described by cryo-EM (Loop-Helix) and a state that was relatively unique to the binding ligands in the monomer (Helix-Helix). In the dimers and trimers, we identified a series of additional more flexible and potential intermediate states that were unique to the protein-protein interactions of the protomers. This analysis was the key for understanding the stability and allostery of these assemblies as well as the overall hexamer. This would have been a challenge, or even impossible, to characterize and understand using the RMSD-based algorithm or CATS, as they provide a surplus of clusters making it exceedingly cumbersome to go through each representative structure individually and to visualize it with VMD. With CATS+, this process would not have been as laborious due to the fewer clusters. However, as mentioned, CATS+ did not even identify some of the important conformations in several of the setups. Both the RMSD-based algorithm and CATS+ rely on visualization with software such as VMD to observe potential structural changes. StELa provides the representative vectors that easily indicate the structure type directly by matching the centroid labels to the appropriate region on the Ramachandran plot. This allows us to be more confident that the structure is more accurately represented, without relying on visualization alone.

### Dihedral angles based clustering algorithms are challenged by the presence of substantial beta-sheet structures

The ability of StELa to correctly cluster structures depends on how well the initial centroid based clustering step separates the regions in the Ramachandran plot, which we found to generally perform better in katanin than in tau. Notably, the fewer centroids used, the less distinct the resulting vectors are going to be. Furthermore, similar to CATS, we also found it challenging to characterize the beta region. The beta region of the Ramachandran plot includes straight beta strands as well as sheets, not distinguishing between the two very different assemblies. StELa has no way of identifying a difference between consecutive straight beta strands and the sheet conformation, which is tertiary in nature, although it does a very good job of characterizing and identifying alpha helices. Still, we found that StELa performs the best at extracting significant structures from the landscape with reasonable populations in comparison to the RMSD-based algorithm and CATS/CATS+.

## Data and Software Availability

The MD trajectory frames for the various systems simulated by the Dima group and tested in this work is available upon request from the corresponding author. The code for StELa can be found on our Github page at https://github.com/DimaUClab/StELa-Protein-Structure-Clustering-Algorithm.

## Supporting information

Supplemental results, tables, figures

## Acknowledgement

We thank Joan Emma Shea and Margaret Cheung for providing the MD simulations files for TAU/R2. This research was funded by the National Science Foundation (NSF) MCB-1817948 (to RID). S.M was supported through the NSF Research Experience for Undergraduates in Chemistry grant CHE-1950244. This work used the Extreme Science and Engineering Discovery Environment (XSEDE) through allocation TG–BIO210094 to RID.

## Supporting Information Available

Additional details of clustering the HBD tip for each protomer in the discussed quaternary assemblies; the equation for the Calinski-Harabasz index used to determine the number of unique clusters; the equation for the silhouette score used to determine the number of unique clusters; the simulation time for the tau systems; the simulation time for the katanin systems; the variance covered by the first two principal components in each of the studied systems; the populations for the top five clusters with each clustering method for each of the R2/Tau systems; the populations for the top five clusters with each clustering method for the R4-R’/Tau system; the populations for the top five clusters with each clustering method for each of the katanin monomer systems; the populations for the top five clusters with each clustering method for each protomer of the katanin ABC trimer systems; the RMSD over time for the R4-R’/Tau simulations; a katanin monomer illustrating the different indicating interfaces and highlighting the HBD tip with a cartoon to explain the lower order oligomers of katanin; the distribution of the phi and psi angle for the COMPLEX setup of the HBD tip of the katanin monomer; the result of KMeans clustering on the Ramachandran plot for the HBD tip of the APO set up of the protomer B in the Ring trimer; the FEL for the HBD tip of the APO set up of protomer C in the Ring trimer expressed in *R_g_*-*R_E_E* space v. PC1-PC2 space; the PC1-PC2 FEL of each R2/Tau system; the PC1-PC2 FEL of each katanin monomer system; the PC1-PC2 FEL of each katanin protomer system; the Calinski-Harabasz score and Silhouette score plots for the R2/Tau in water with the indicated local maxiumums; the dendrogram for the R2/Tau in water; the FEL with indicated cluster centers for each clustering method with representative structures for each indicated FEL minima for R2/Tau in TMAO; the FEL with indicated cluster centers for each clustering method with representative structures for each indicated FEL minima for R2/Tau in UREA; the FEL with indicated cluster centers for each clustering method with representative structures for each indicated FEL minima for katanin monomer in the NUCLEOTIDE setup; the FEL with indicated cluster centers for each clustering method with representative structures for each indicated FEL minima for katanin monomer in the SUB setup; the FEL with indicated cluster centers for each clustering method with representative structures for each indicated FEL minima for katanin monomer in the APO setup; the FEL with indicated cluster centers for each clustering method with representative structures for each indicated FEL minima for protomer A from the katanin ABC trimer in the COMPLEX setup; the FEL with indicated cluster centers for each clustering method with representative structures for each indicated FEL minima for protomer A from the katanin ABC trimer in the APO setup; the FEL with indicated cluster centers for each clustering method with representative structures for each indicated FEL minima for protomer B from the katanin ABC trimer in the COMPLEX setup; the FEL with indicated cluster centers for each clustering method with representative structures for each indicated FEL minima for protomer C from the katanin ABC trimer in the COMPLEX setup; the FEL with indicated cluster centers for each clustering method with representative structures for each indicated FEL minima for protomer C from the katanin ABC trimer in the APO setup; the RMSD distributions for the R2/Tau systems and for the katanin monomer systems; the Ramachandran plot for R2/Tau in water and the HBD tip of the monomer from the COMPLEX setup.

