## Supplemental results, tables, figures for "Searching for Structure: Characterizing the Protein Conformational Landscape with Clustering-based Algorithms"

### Additional Results

#### Protein Fragment in Quaternary Assemblies: the HBD tip of each Protomer of the Katanin Ring Trimer

In the COMPLEX state, protomer A identified four distinct regions in the FEL with high barriers between them and a clear separation of region (ii) from the rest as seen in **Figure S16**. Region (iii) was the only one to capture the N-terminal loop to helix transition while the others identify different orientations of the loop region and lengths of the C-terminal helix. Protomer B was found to sample the space in a very similar way, as seen in **Figure S18**, identifying similar regions with the addition of a more shallow region connecting the two disjointed areas of space as the result of the additional component of the convex interface. The structures from the identified regions represent similar types of structures as found for protomer A, however the region associated with the loop to helix transition (iii) is more separate from the others. The FEL of protomer C was found to experience more shallow barriers between six different sampled regions as seen in **Figure S19**. All of the sampled regions represent variations with the N-terminal loop to helix transition except for region (ii). StELa did the best at capturing clusters from each of the regions from the described protomers. There were two more shallow minima-like regions in protomer C that were not identified by any of the three methods. CATS+ sampled three out of four regions in protomer A, three out of five regions in protomer B, and only one out of five regions in protomer C. The RMSD-based algorithm only identified one in protomer A and two in protomers B and C.

In general, the removal of the cofactors results in fewer regions. Protomer A, the protomer that is missing the nucleotide in the COMPLEX setup and is the protomer observed to dissociate in the PC motions, is observed to go from four to three regions.[1] In the APO setup, shown in **Figure S17**, the structures associated with the corresponding regions are distinct. Region (i) corresponds to the loop to helix transition, region (ii) corresponds to a more flexible state, and region (iii) corresponds to a slightly longer C-terminal helix. Protomer B, discussed in the main text, experiences a dramatic shift in the FEL in the absence of the cofactors as seen in **Figure 4**. The identified structures are found to represent generally more flexible configurations. This is likely a requirement of the described global changes in protomer B to account for the motions of A and C. In protomer C, the removal of the cofactors results in two deeper minima and two more shallow regions as shown in **Figure S20**. The structures associated with these regions are found to be even more flexible than those from the COMPLEX setup. StELa was able to capture and characterize each of the regions of interest across the protomers in the APO setup. CATS+ was able to identify three out of three regions in protomer A, three out of five regions in protomer B and two out of four regions in promoter C. The RMSD-based algorithm identified one region in protomers A and B, and two regions in protomer C. As the analysis of the PC motions from our previous study indicated that protomer A will dissociate from the ABC-trimer due to the increase in disorder of the NBD and protomer B will remain bound to protomer C, the comparison of the HBD fragment for protomers A and B is particularly interesting. In order for B to stay attached to C, the region is required to lower the conformational energy barriers, allowing it to become more flexible. In comparison, A is found to have higher energy barriers with fewer conformations, characterizing a relatively more rigid landscape.

### Additional Methods

The included equations below were used to calculate the scores for identifying the appropriate number of clusters for the datasets used in StELa as discussed in the *Results*.

**Equation S1**: Calculating the Calinski-Harabasz Index ($s$) on a dataset $D = [d_1, d_2, \ldots d_N]$ of size $n_D$ that has been clustered into $k$ clusters [2,3]:

$$s = \frac{tr(B_k)}{tr(W_k)} \times \frac{n_D - k}{k - 1} \quad (1)$$

$$B_k = \sum_{q=1}^{k} n_q (c_q - c_D)(c_q - c_D)^T \quad (2)$$

$$W_k = \sum_{q=1}^{k} \sum_{1 \epsilon C_q} (x - c_q)(x - c_q)^T \quad (3)$$

**Equation S2**: Calculating the Silhouette Score ($s$) where a is the mean intra-cluster distance and b is the mean nearest-cluster distance [4,3]:

$$s(i) = \frac{b(i) - a(i)}{max\,\{a(i), b(i)\}}$$

**Table S1**: Table of the tau simulation time. All R2/Tau trajectories are sourced from the Shea group.[5]

| System | Trajectory (Total Time) |
|---|---|
| R2/Tau: Water | 1 (120ns) |
| R2/Tau: TMAO | 1 (200ns) |
| R2/Tau: Urea | 1 (150ns) |
| R4-R'/Tau | 4 (800ns) |

**Table S2**: Table of the simulation time used in this study, taken from our previous study of lower order oligomers.[1]

| Katanin | Trajectory (Time) | Total Simulation Time |
|---|---|---|
| Monomer/COMPLEX | 3 (50ns) | 150 ns |
| Monomer/APO | 3 (50ns) | 150 ns |
| ABC/COMPLEX | 4 (100ns) | 400ns |
| ABC/APO | 4 (100ns) | 400ns |

**Table S3**: Table of the contribution for the first two PCs of each indicated system.

| | R2/Tau: **Water** | R2/Tau: **TMAO** | R2/Tau: **UREA** | R4-R'/Tau: **Water** |
|---|---|---|---|---|
| PC1 | 23.1% | 22.8% | 18.0% | 27.4% |
| PC2 | 8.6% | 9.9% | 14.6% | 14.4% |
| Total | 31.7% | 32.7% | 32.6% | 41.8% |

| Katanin HBD tip | Monomer **COMPLEX** | Monomer **NUC** | Monomer **SUB** | Monomer **APO** |
|---|---|---|---|---|
| PC1 | 49.6% | 26.6% | 44.8% | 38.2% |
| PC2 | 17.9% | 18.8% | 16.3% | 15.0% |
| Total | 67.5% | 45.4% | 61.1% | 53.2% |

| Katanin HBD tip | Trimer - A | Trimer - B | Trimer - C |
|---|---|---|---|
| | **COMPLEX** | **COMPLEX** | **COMPLEX** |
| PC1 | 52.6% | 34.9% | 32.5% |
| PC2 | 11.0% | 21.6% | 22.0% |
| Total | 63.6% | 56.5 | 54.5 |
| | **APO** | **APO** | **APO** |
| PC1 | 40.2% | 30.8% | 33.4% |
| PC2 | 23.9% | 21.0% | 15.5% |
| | 64.1% | 51.8% | 48.9% |

**Table S4**: Table of the top 5 populated clusters identified with each method of each environment of the R2 fragment of Tau. The number of clusters used are indicated in the column header.

| **Water** | GROMOS (1540) | CATS+ (5) | StELa (21) |
|---|---|---|---|
| Cluster 1 | 4.1% | 85.1% | 28.2% |
| Cluster 2 | 4.1% | 9.5% | 12.4% |
| Cluster 3 | 2.1% | 4.8% | 9.1% |
| Cluster 4 | 2.0% | 0.3% | 8.1% |
| Cluster 5 | 1.3% | 0.2% | 6.0% |

| **TMAO** | GROMOS (1928) | CATS+ (10) | StELa (16) |
|---|---|---|---|
| Cluster 1 | 5.6% | 92.9% | 26.6% |
| Cluster 2 | 2.7% | 2.1% | 15.9% |
| Cluster 3 | 2.5% | 1.1% | 12.3% |
| Cluster 4 | 2.3% | 1.0% | 10.4% |
| Cluster 5 | 1.3% | 0.9% | 7.9% |

| **Urea** | GROMOS (2186) | CATS+ (5) | StELa (22) |
|---|---|---|---|
| Cluster 1 | 1.7% | 89.0% | 25.9% |
| Cluster 2 | 1.3% | 7.4% | 13.5% |
| Cluster 3 | 1.1% | 1.9% | 9.4% |
| Cluster 4 | 0.8% | 0.9% | 8.4% |
| Cluster 5 | 0.6% | 0.8% | 8.3% |

**Table S5**: Table of the top 5 populated clusters identified with each method of the R4-R' fragment of Tau. The number of clusters used are indicated in the column header.

| | GROMOS (7203) | CATS+ (1) | StELa (23) |
|---|---|---|---|
| Cluster 1 | 8.3% | 100% | 21.0% |
| Cluster 2 | 1.6% | - | 14.9% |
| Cluster 3 | 1.2% | - | 9.6% |
| Cluster 4 | 0.9% | - | 9.4% |
| Cluster 5 | 0.7% | - | 6.9% |

**Table S6**: Table of the top 5 populated clusters identified with each method of each protomer of Katanin monomer in the COMPLEX, NUC, SUB, & APO setups. The number of clusters used are indicated in the column header.

| COMPLEX **MONOMER** | GROMOS (23) | CATS+ (3) | StELa (18) |
|---|---|---|---|
| Cluster 1 | 38.0% | 55.7% | 30.1% |
| Cluster 2 | 25.5% | 26.5% | 20.1% |
| Cluster 3 | 14.8% | 17.7% | 18.0% |
| Cluster 4 | 6.7% | - | 12.9% |
| Cluster 5 | 5.6% | - | 9.2% |

| NUCLEOTIDE **MONOMER** | GROMOS (37) | CATS+ (6) | StELa (14) |
|---|---|---|---|
| Cluster 1 | 32.2% | 59.1% | 30.3% |
| Cluster 2 | 17.7% | 32.1% | 14.3% |
| Cluster 3 | 10.5% | 4.4% | 13.2% |
| Cluster 4 | 7.6% | 2.3% | 13.1% |
| Cluster 5 | 4.1% | 1.3% | 8.6% |

| SUBSTRATE **MONOMER** | GROMOS (33) | CATS+ (6) | StELa (8) |
|---|---|---|---|
| Cluster 1 | 29.6% | 58.5% | 24.6% |
| Cluster 2 | 22.0% | 22.9% | 16.6% |
| Cluster 3 | 17.6% | 12.8% | 15.5% |
| Cluster 4 | 5.1% | 2.7% | 11.0% |
| Cluster 5 | 3.8% | 1.7% | 10.1% |

| APO **MONOMER** | GROMOS (37) | CATS+ (5) | StELa (14) |
|---|---|---|---|
| Cluster 1 | 26.6% | 57.1% | 38.7% |
| Cluster 2 | 15.3% | 23.4% | 18.7% |
| Cluster 3 | 10.1% | 16.5% | 17.0% |
| Cluster 4 | 8.1% | 1.6% | 6.7% |
| Cluster 5 | 7.7% | 1.5% | 6.4% |

**Table S7**: Table of the top 5 populated clusters identified with each method of each protomer of Katanin RING ABC trimer in the COMPLEX & APO setups. The number of clusters used are indicated in the column header.

| 6UGE-ABC-COMP **Protomer A** | GROMOS (21) | CATS+ (12) | StELa (12) |
|---|---|---|---|
| Cluster 1 | 24.6% | 48.8% | 41.7% |
| Cluster 2 | 23.6% | 12.0% | 20.8% |
| Cluster 3 | 21.2% | 9.6% | 13.7% |
| Cluster 4 | 19.9% | 6.3% | 8.1% |
| Cluster 5 | 3.5% | 5.3% | 6.5% |

| 6UGE-ABC-COMP **Protomer B** | GROMOS (17) | CATS+ (8) | StELa (13) |
|---|---|---|---|
| Cluster 1 | 49.5% | 60.5% | 38.4% |
| Cluster 2 | 20.2% | 22.3% | 28.0% |
| Cluster 3 | 16.4% | 5.3% | 14.7% |
| Cluster 4 | 4.1% | 3.8% | 7.1% |
| Cluster 5 | 3.6% | 3.0% | 3.1% |

| 6UGE-ABC-COMP **Protomer C** | GROMOS (87) | CATS+ (4) | StELa (16) |
|---|---|---|---|
| Cluster 1 | 24.2% | 79.0% | 25.1% |
| Cluster 2 | 9.7% | 13.1% | 18.2% |
| Cluster 3 | 9.1% | 4.8% | 11.2% |
| Cluster 4 | 8.5% | 3.2% | 9.2% |
| Cluster 5 | 7.5% | - | 7.1% |

| 6UGE-ABC-APO **Protomer A** | GROMOS (25) | CATS+ (8) | StELa (8) |
|---|---|---|---|
| Cluster 1 | 41.0% | 61.1% | 45.7% |
| Cluster 2 | 26.4% | 13.9% | 26.8% |
| Cluster 3 | 20.0% | 6.8% | 14.4% |
| Cluster 4 | 3.6% | 6.0% | 11.3% |
| Cluster 5 | 2.0% | 4.8% | 0.9% |

| 6UGE-ABC-APO **Protomer B** | GROMOS (25) | CATS+ (10) | StELa (17) |
|---|---|---|---|
| Cluster 1 | 31.3% | 39.3% | 20.5% |
| Cluster 2 | 25.8% | 20.8% | 18.0% |
| Cluster 3 | 15.35% | 11.0% | 13.2% |
| Cluster 4 | 6.5% | 9.3% | 10.4% |
| Cluster 5 | 5.9% | 5.7% | 10.1% |

| 6UGE-ABC-APO **Protomer C** | GROMOS (68) | CATS+ (8) | StELa (17) |
|---|---|---|---|
| Cluster 1 | 43.5% | 54.1% | 40.0% |
| Cluster 2 | 15.3% | 15.5% | 20.7% |
| Cluster 3 | 7.7% | 10.0% | 11.3% |
| Cluster 4 | 4.2% | 5.9% | 9.3% |
| Cluster 5 | 3.7% | 5.8% | 7.5% |

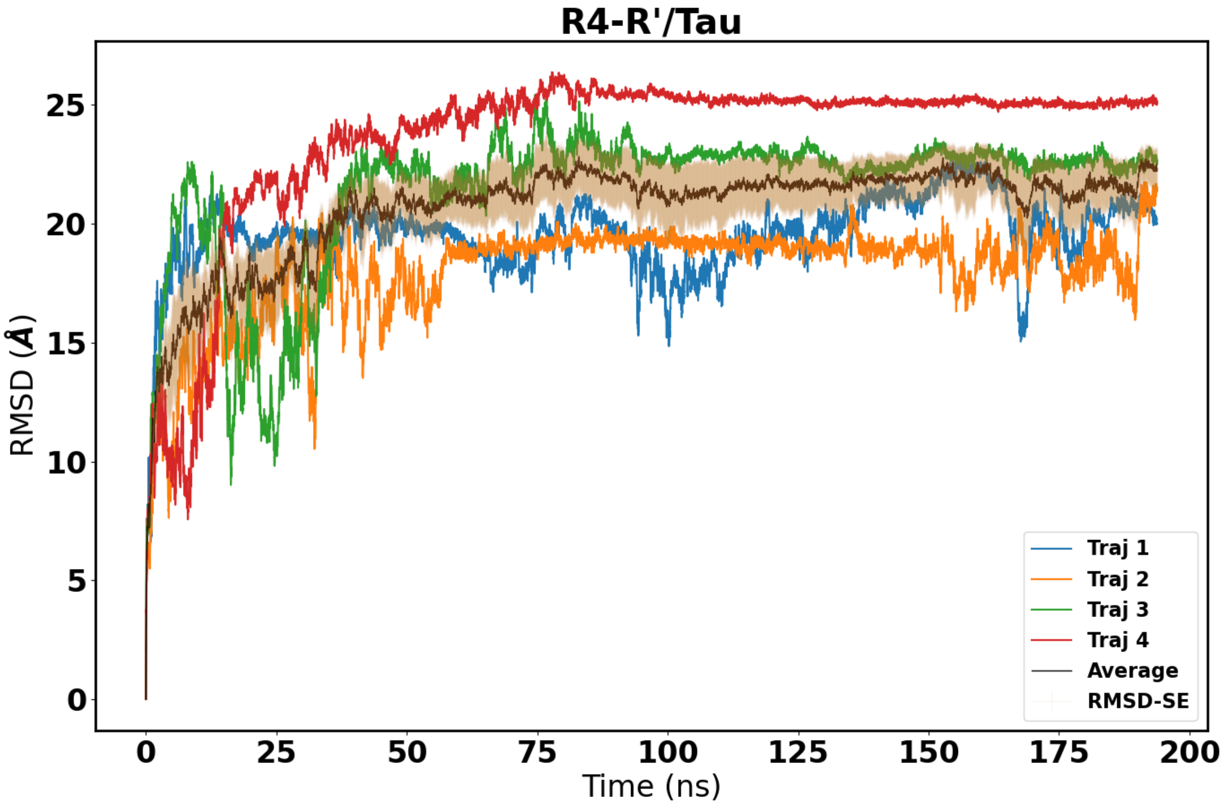


**Figure S1**: Global RMSD of R4-R'/Tau backbone atoms. Each trajectory is in a different color as indicated in the legend, and the average across all trajectories is shown in black with associated error bars in tan. Average error is 1.179.

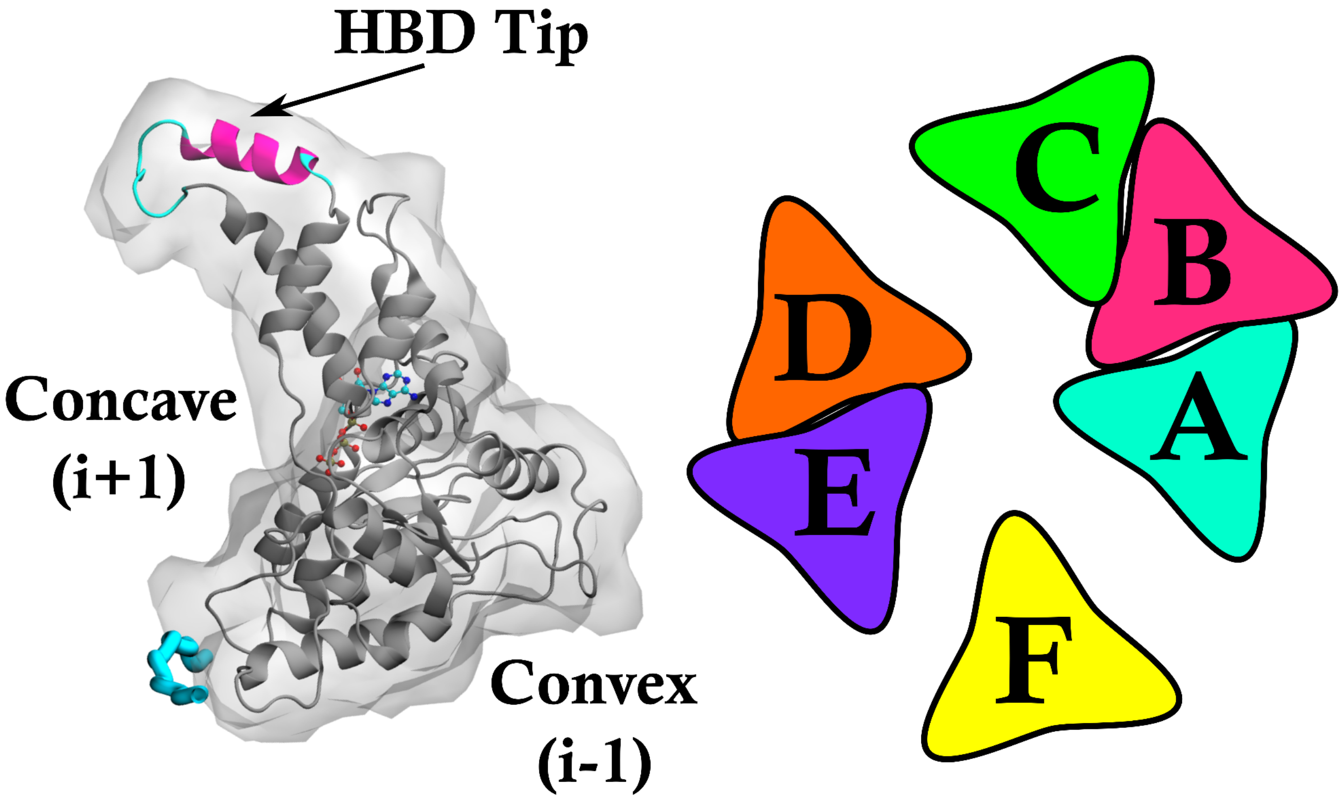


**Figure S2**: The cartoon on the right shows how the hexamer of Katanin was broken up into lower order oligomers in our previous study.[1] For katanin, we carried out simulations of the ABC trimer, the AB and BC dimer, and the Monomer, which we took protomer A. The location of the interfaces (concave & convex) is indicated on the shown monomer made with VMD (left).[6] The ATP is shown in the binding pocket (cpk) and the polyglutamate substrate is indicated in light blue. In addition, we indicate where the "HBD tip", the subject of our clustering analysis, is located in the protomer.

a.)

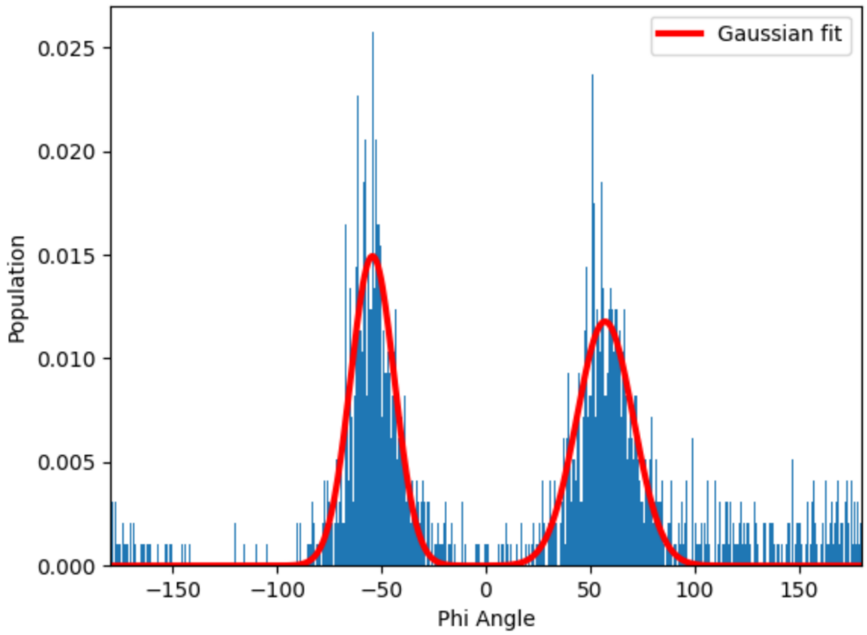


b.)

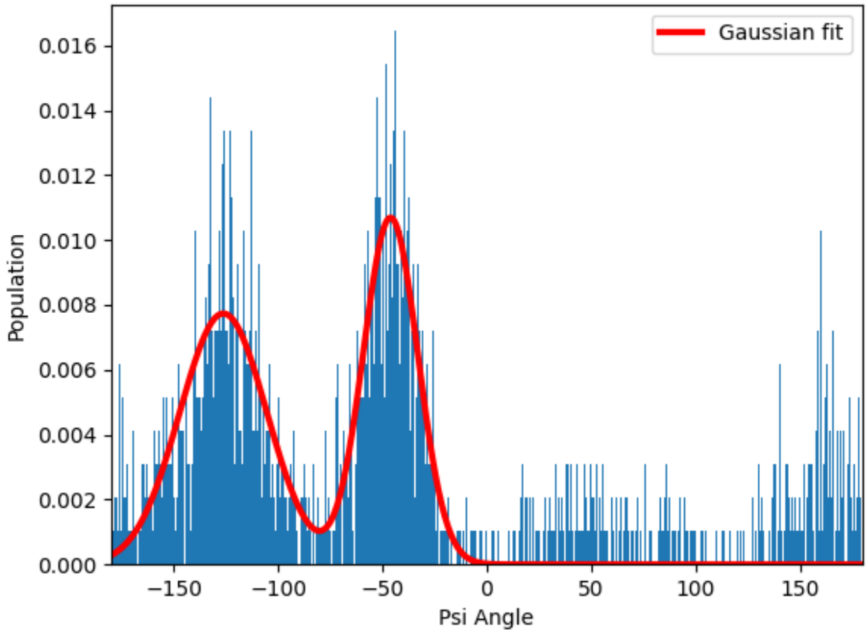


**Figure S3**: The distribution of the phi (a) and psi (b) angles between R422 & G423 for the HBD tip of the katanin monomer in the presence of both cofactors (COMPLEX).[1,7]

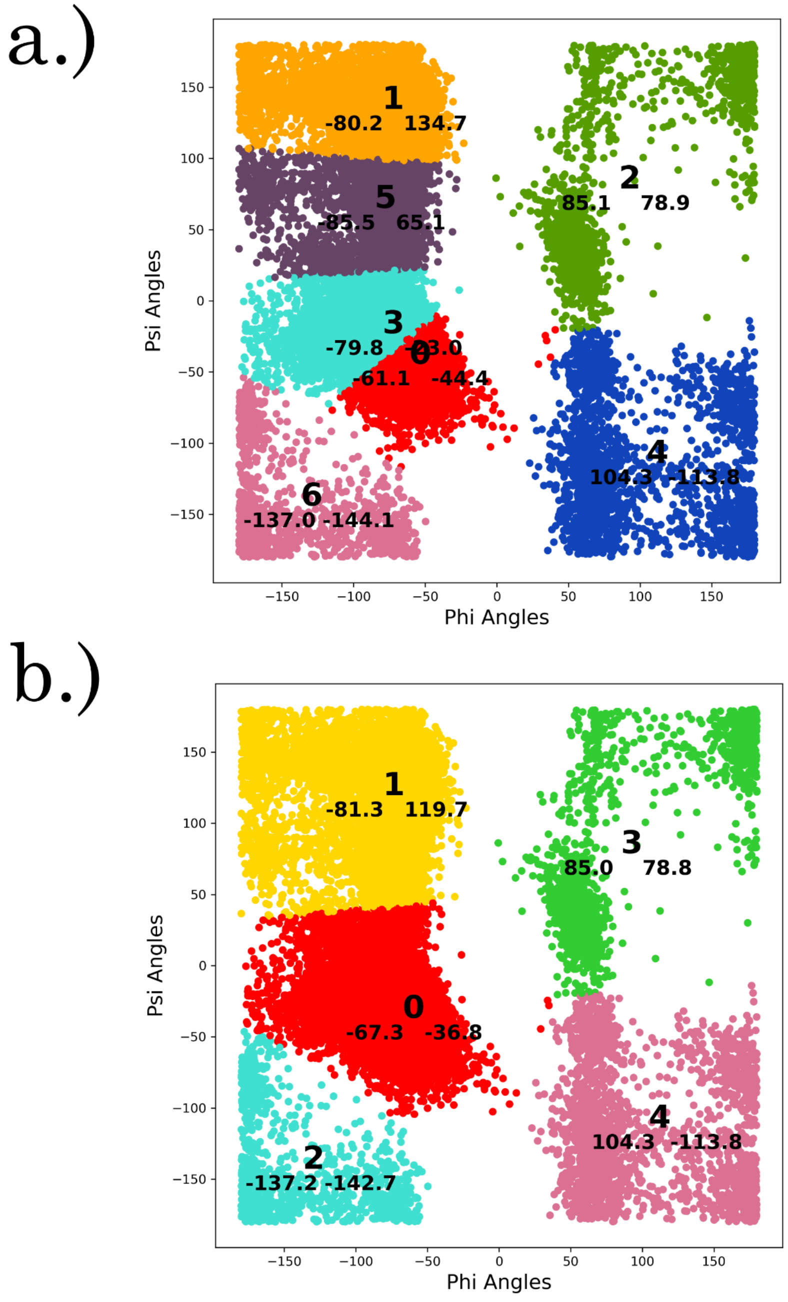


**Figure S4**: The result of using centroid based clustering (K-Means) on the Ramachandran plot, colored according to the cluster IDs, for the "HBD tip" fragment of Protomer B from the Ring conformation in the APO state. (a) When using 7 clusters, the region associated with Alpha helices was split which made characterizing or sorting through the vectors more difficult and therefore, (b) 5 clusters was used for characterizing the structure vectors for this example - even though 7 clusters is a more descriptive way to characterize the differences between the regions.

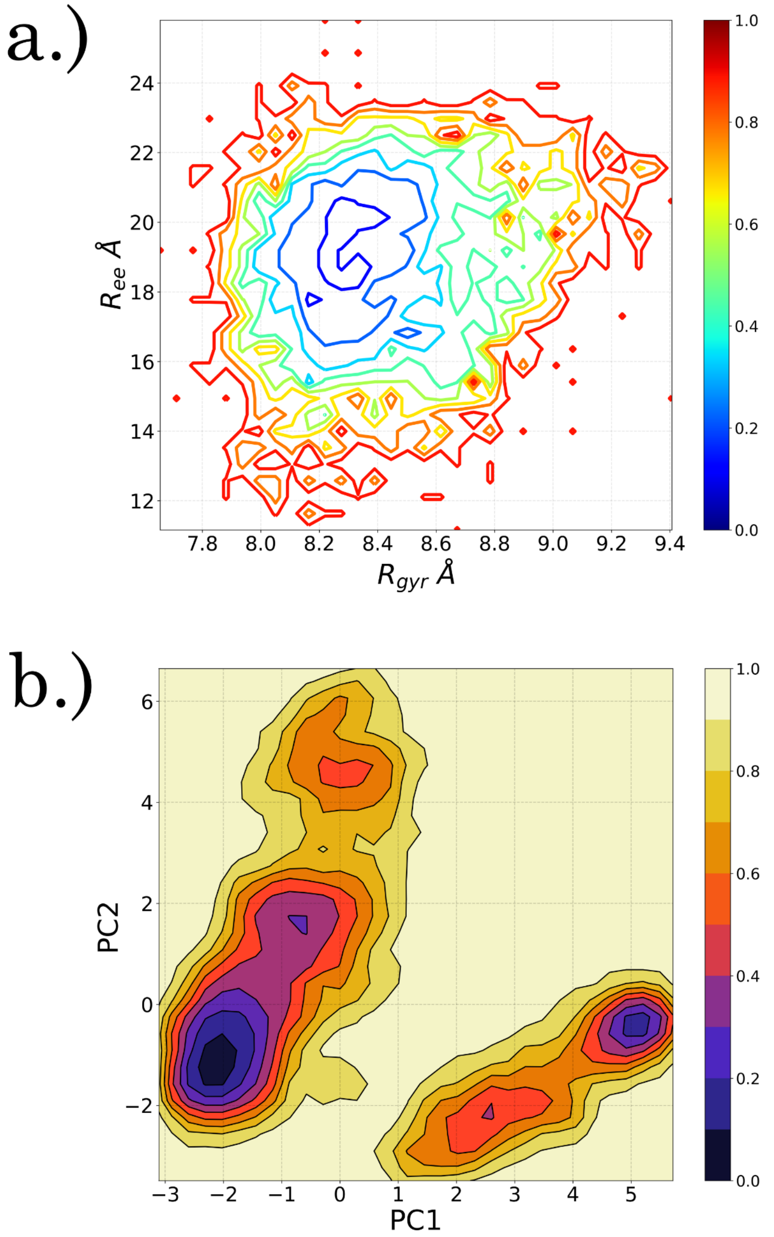


**Figure S5**: The free energy landscape for the "HBD tip" fragment from Protomer C in the Ring-ABC-APO set up in RGYR v. REE space. As seen here, this space only describes a single minima in the conformational landscape. Based on the FEL expressed in PC space,[8] there are 2 very distinct regions, 2 deep minimas of interest and 2 additional small and shallow regions of potential interest (also shown in **Figure S20**). The RGYR v. REE space does not provide adequate detail for describing the conformational space of the protein fragments presented in this study, although it does a better job with more behaved systems like katanin rather than IDPs.[1,7]

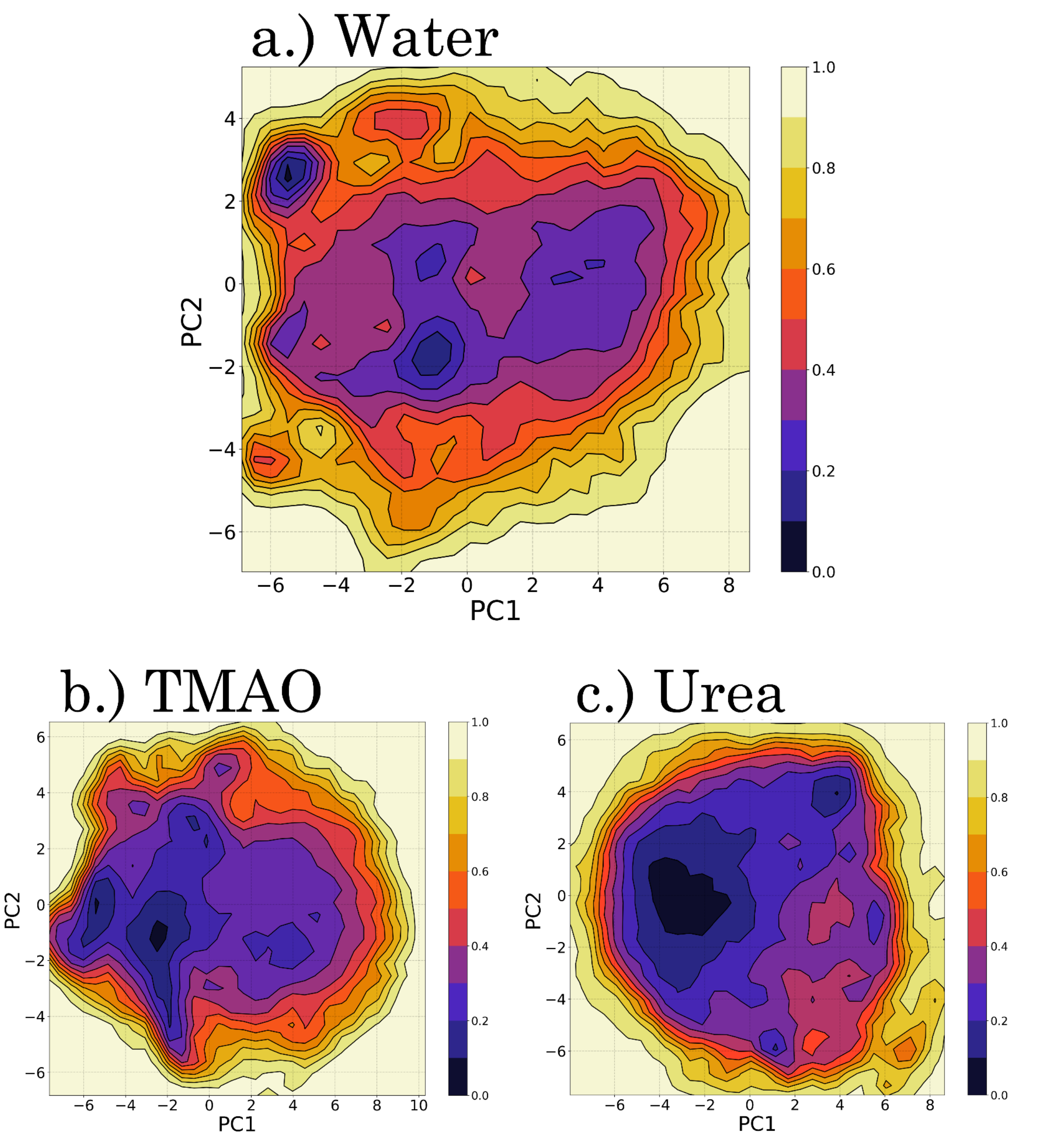


**Figure S6:** The Free Energy Landscapes of the R2 fragment of Tau in PC1/PC2 space for each solvent included in this study: (a) Water, (b) TMAO, & (c) Urea.[5,7]

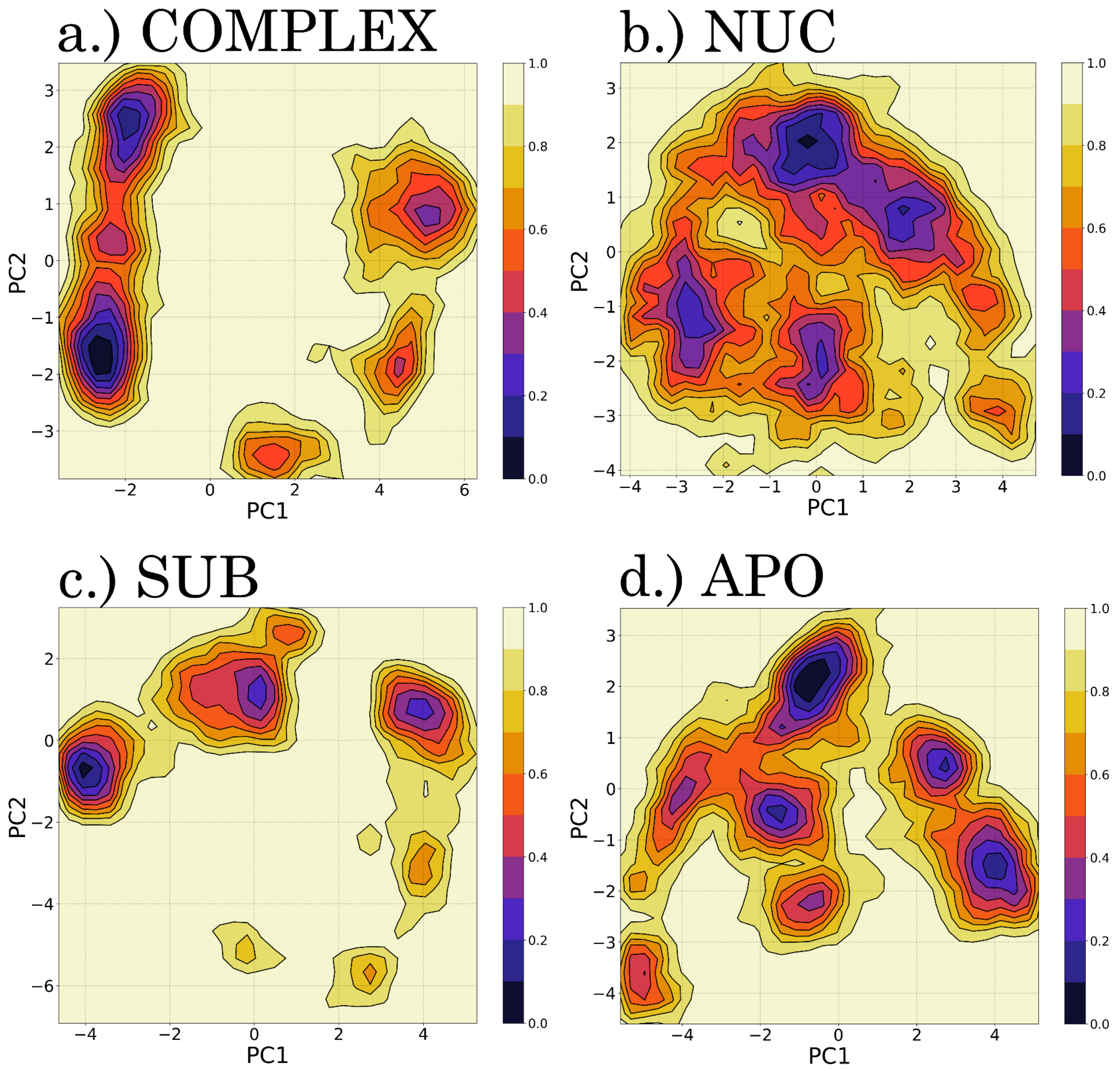


**Figure S7:** The Free Energy Landscapes of the HBD tip fragment from the Monomer in the presence of the cofactors (ATP and the minimal MT substrate) in PC1/PC2 space: (a) COMPLEX, (b) NUC, (c) SUB, (d) APO.[1,7]

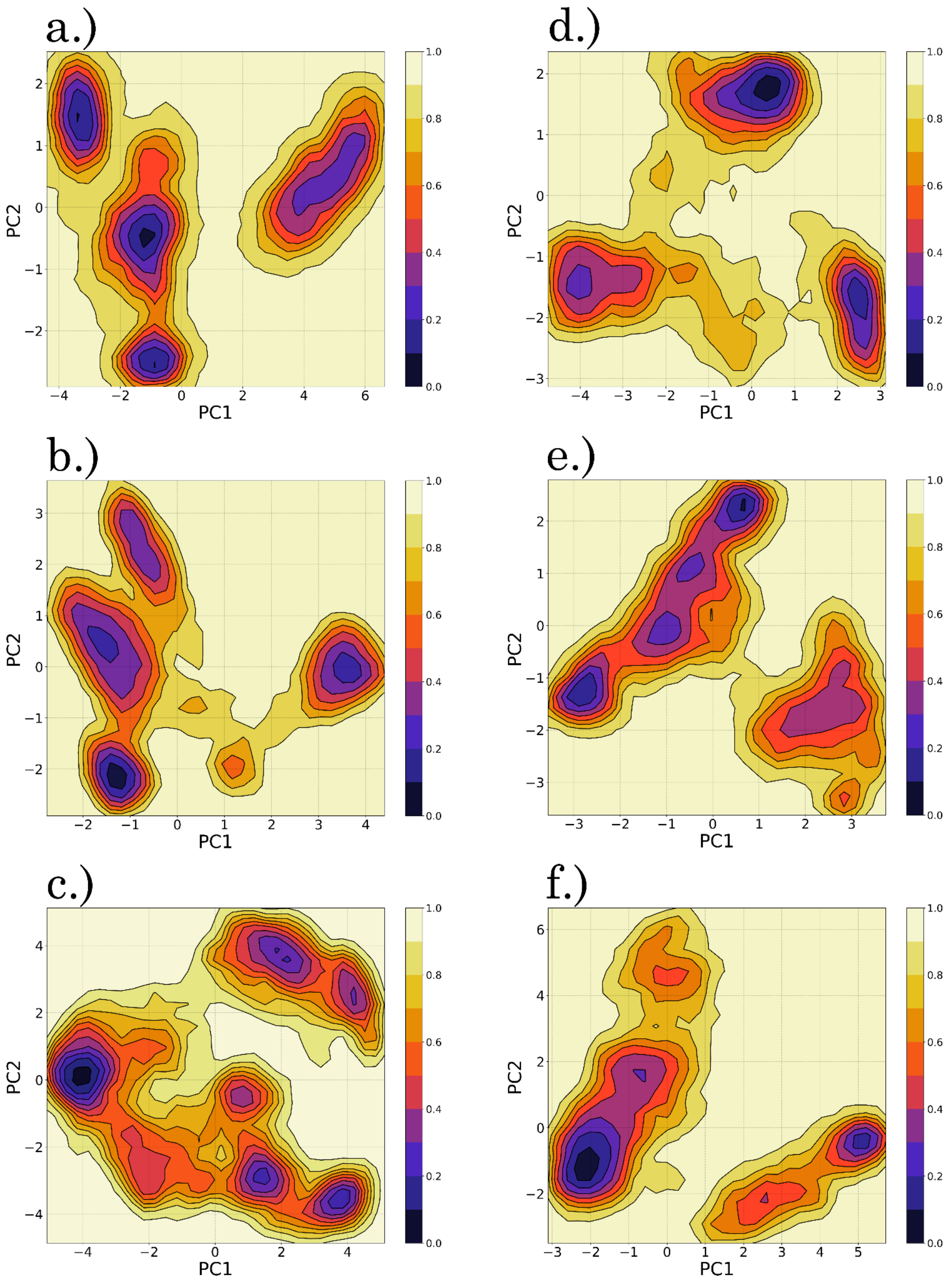


**Figure S8:** The Free Energy Landscapes of the HBD tip fragment of each protomer in the ABC trimer from the Ring conformation in the presence of the ligands ((a-c) COMPLEX, (d-f) APO) in PC1/PC2 space for each protomer included in this study: (a,d) Protomer A, (b,e) Protomer B, (c,f) Protomer C. It is important to note that in the COMPLEX state, the nucleotide is missing from Protomer A in the ring conformation.[1,7]

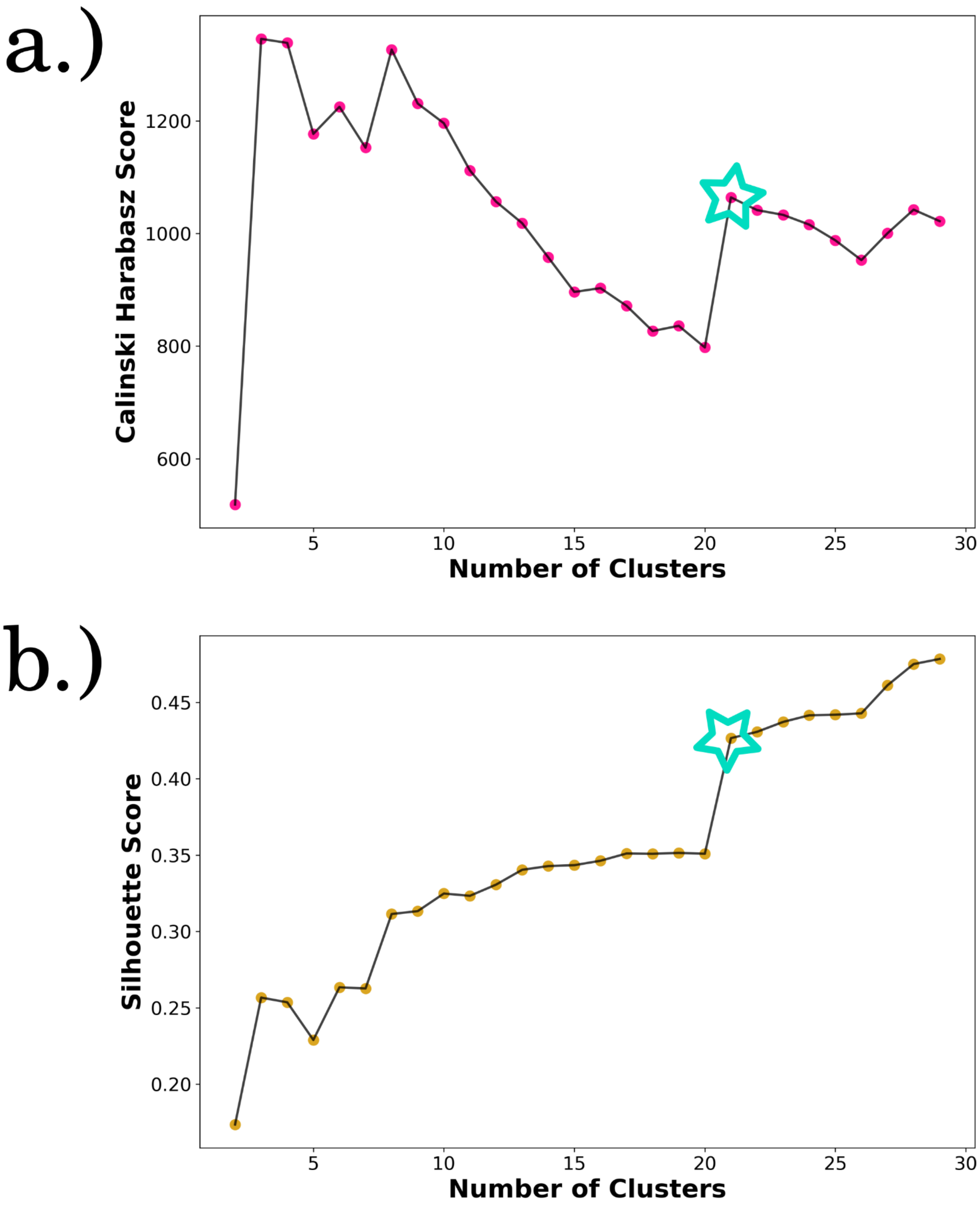


**Figure S9:** The calculated the (a.) Calinski-Harabasz[2] and (b.) Silhouette scores[4,9] plotted against the number of clusters taken for R2/Tau in water.[3] The local maximums, indicated with stars, suggest an optimal number of clusters for this set up being 21.

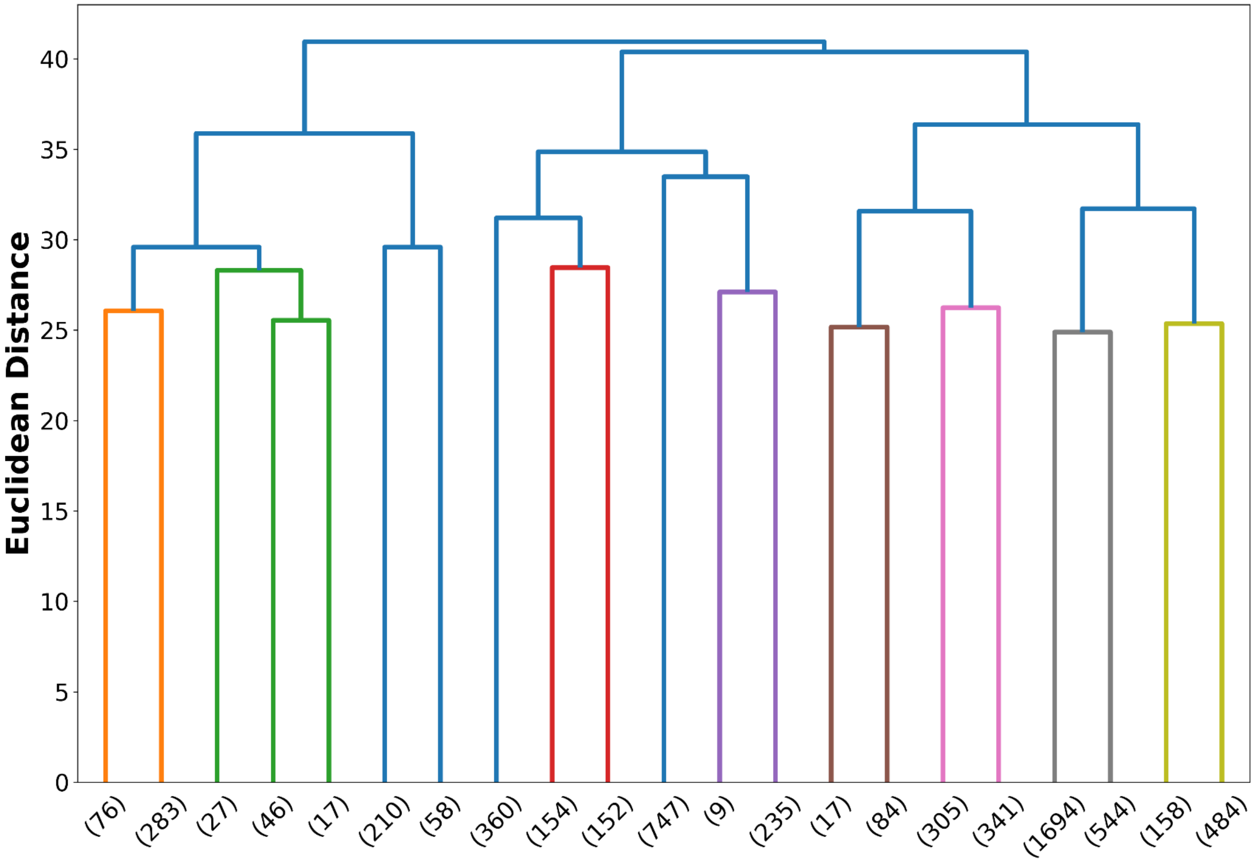


**Figure S10:** The dendrogram from utilizing complete linkage hierarchical clustering to cluster the representative vectors from R2/Tau in water.[3] The number of clusters taken was 21 for this system which is expressed at a euclidean distance of 25. The visualization of this example is truncated to the number of clusters taken so the resulting number of data points are indicated at the branch tick.

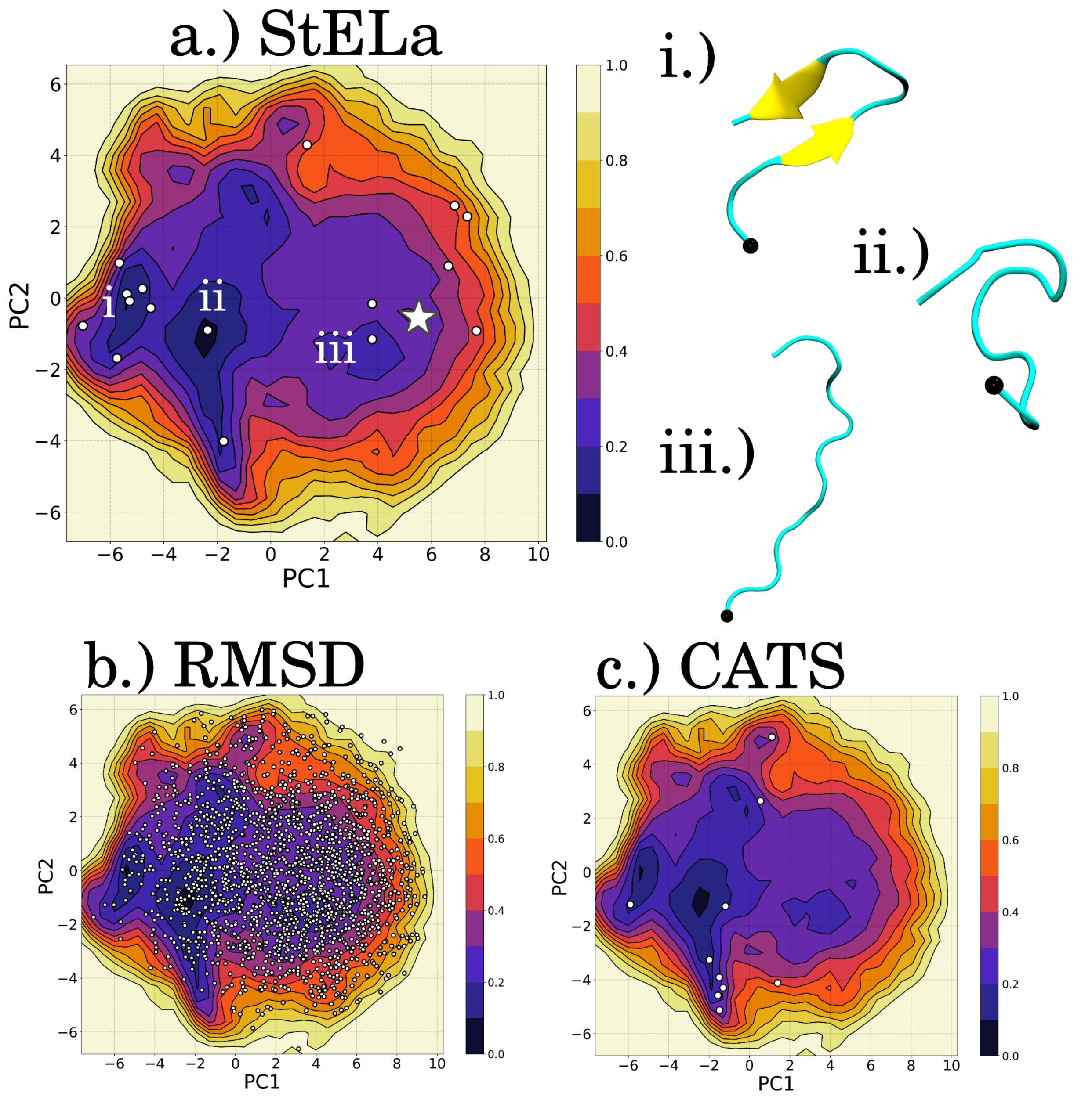


**Figure S11**: The representative structures, made with VMD,[6] for the identified clusters for the R2 fragment of **Tau** in **TMAO** are plotted on the FEL in PC1/PC2 space for each method: (a) StELa, (b) RMSD and (c) CATS+.[1,7] The white star in (a) indicates where the starting structure is found in the beginning of the simulation. Representative structures (i-iii) were extracted to represent the minima indicated in (a). The N-terminal end of the structure is indicated with a black bead.

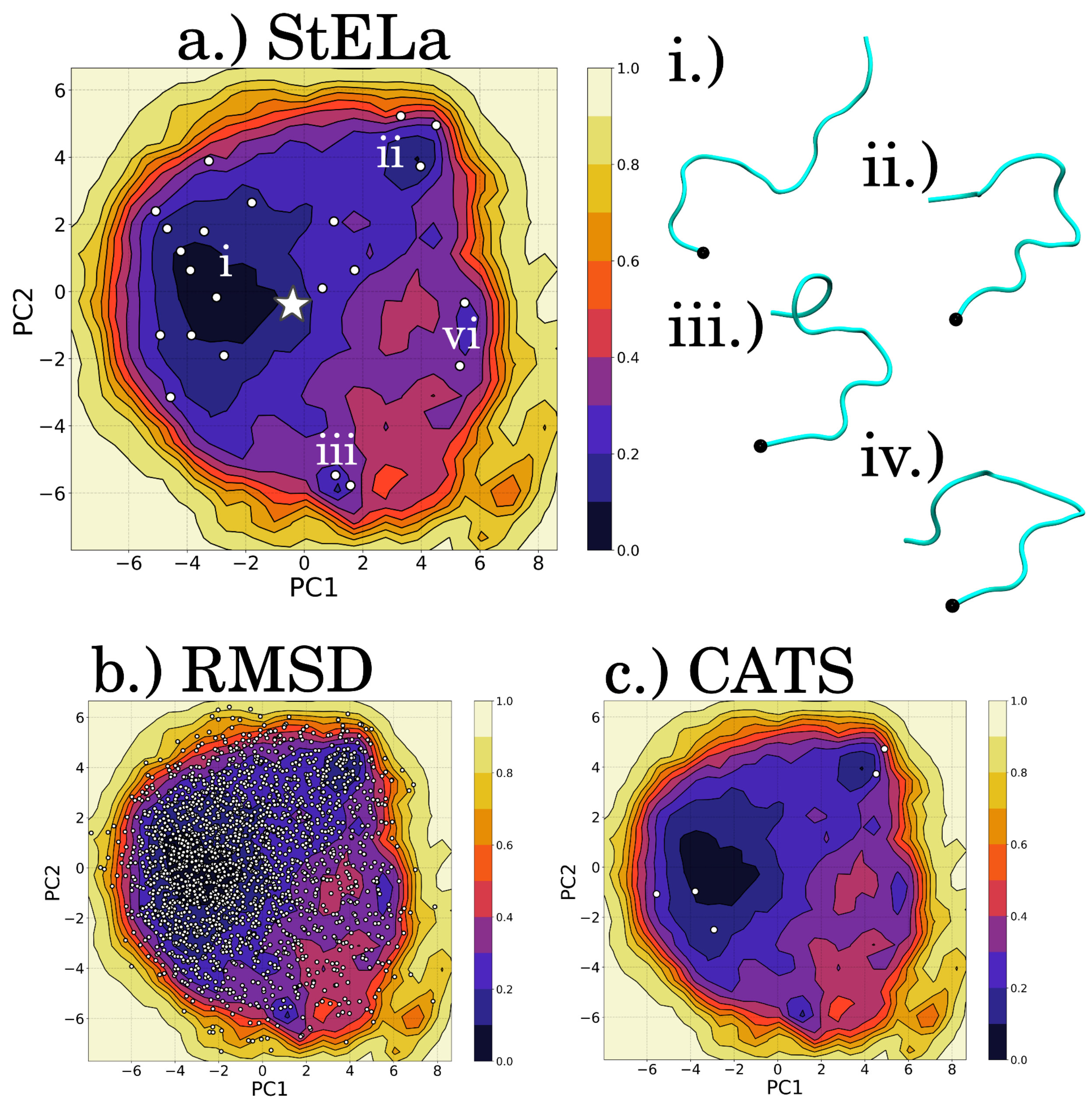


**Figure S12**: The representative structures, made with VMD,[6] for the identified clusters for the R2 fragment of **Tau** in **Urea** are plotted on the FEL in PC1/PC2 space for each method: (a) StELa, (b) RMSD and (c) CATS+.[1,7] The white star in (a) indicates where the starting structure is found in the beginning of the simulation. Representative structures (i-iv) were extracted to represent the minima indicated in (a). The N-terminal end of the structure is indicated with a black bead.

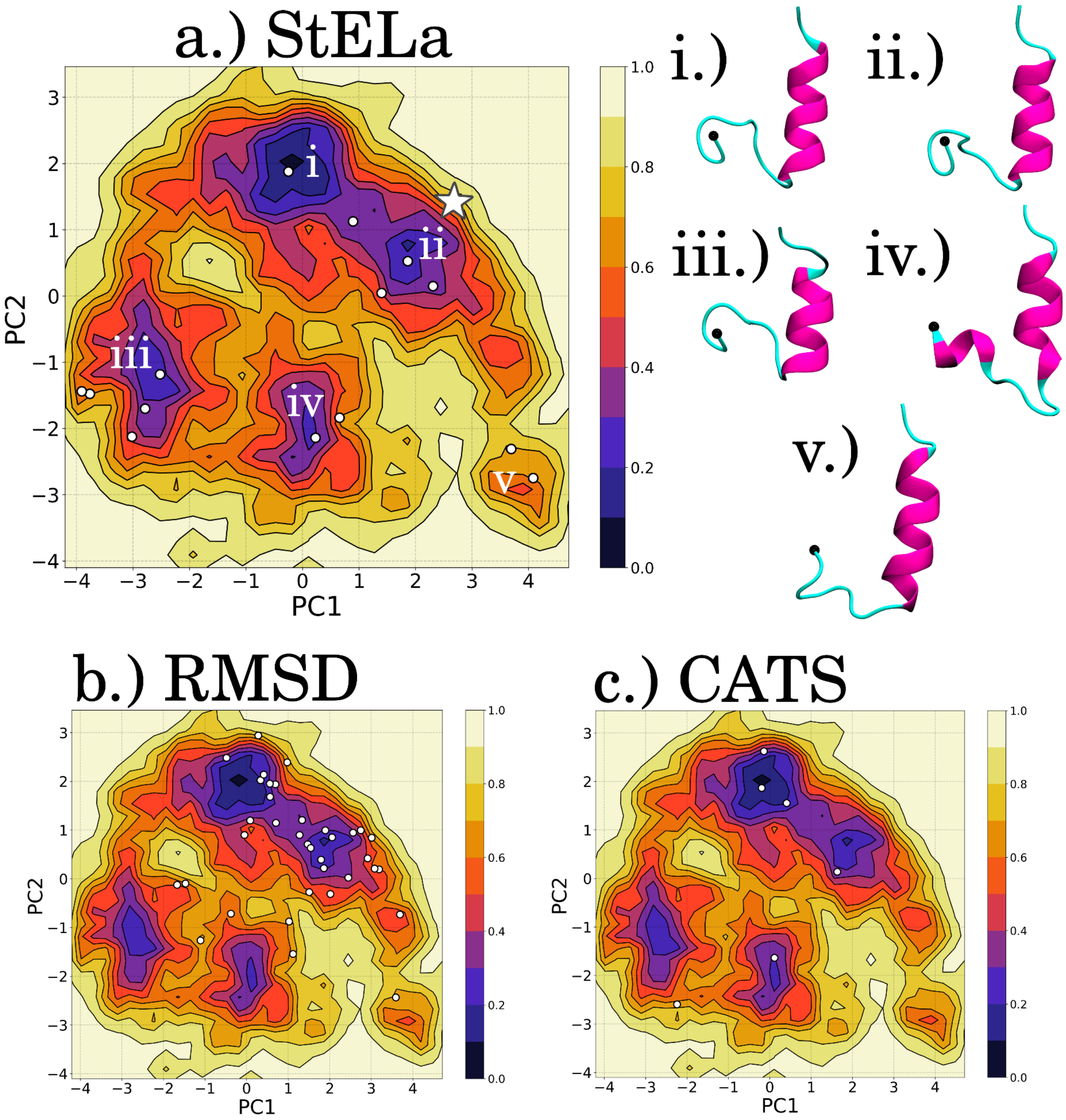


**Figure S13**: The representative structures, made with VMD,[6] for the identified clusters for the HBD tip of the **Monomer NUC** set up are plotted on the FEL in PC1/PC2 space for each method: (a) StELa, (b) RMSD and (c) CATS+.[1,7] The white star in (a) indicates where the starting structure is found in the beginning of the simulation. Interestingly, region iv is associated with the Loop to Helix transition associated with the presence of the cofactors in our previous study.[1] Representative structures (i-v) were extracted to represent the minima indicated in (a). The N-terminal end of the structure is indicated with a black bead.

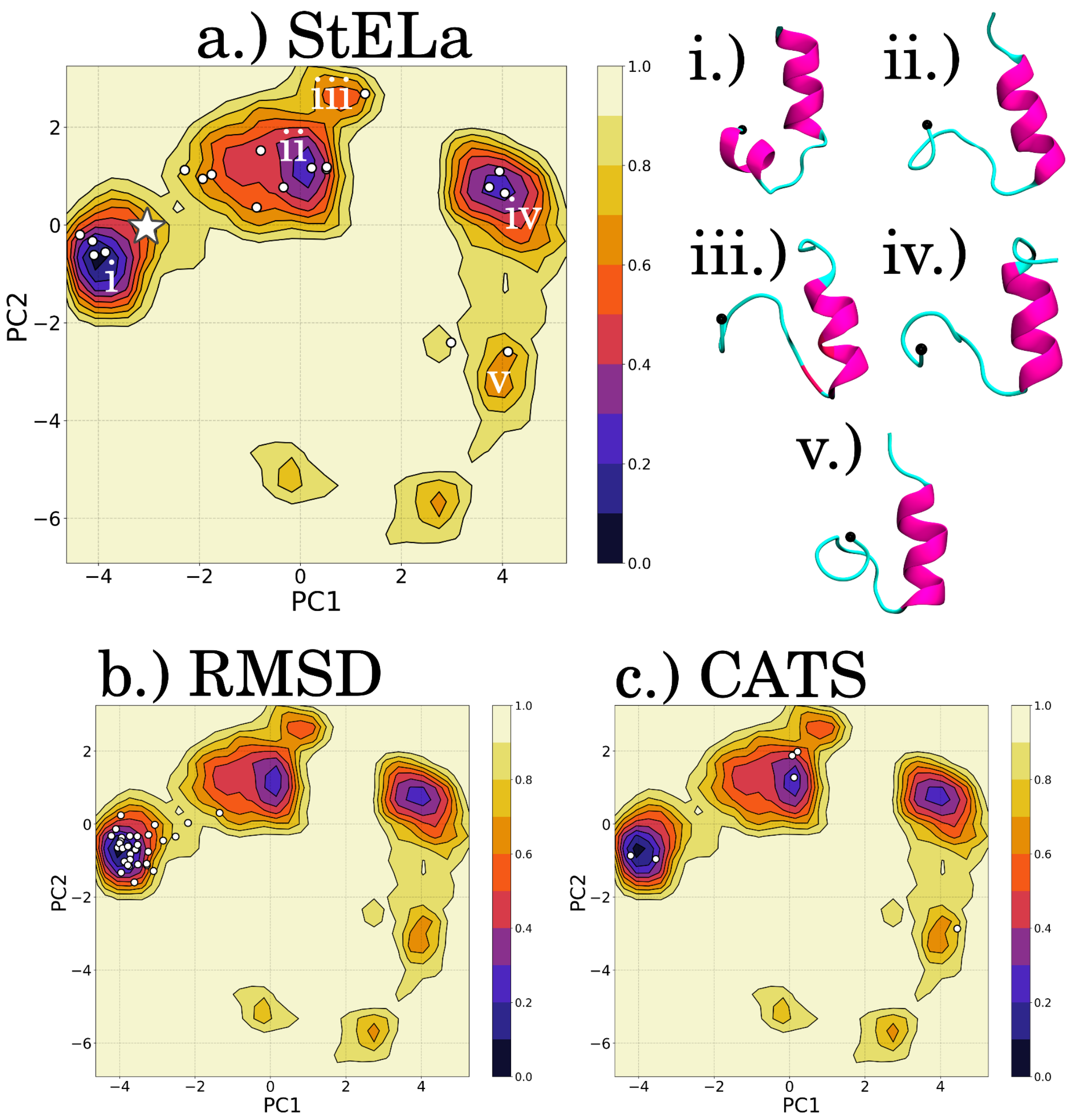


**Figure S14:** The representative structures, made with VMD,[6] for the identified clusters for the HBD tip of the **Monomer SUB** set up are plotted on the FEL in PC1/PC2 space for each method: (a) StELa, (b) RMSD and (c) CATS+.[1,7] The white star in (a) indicates where the starting structure is found in the beginning of the simulation. Representative structures (i-v) were extracted to represent the minima indicated in (a). The N-terminal end of the structure is indicated with a black bead. Neither StELa nor CATS+ identified a cluster in two of the higher energy regions. Interestingly, region i is associated with the Loop to Helix transition associated with the presence of the cofactors (more prominent in the presence of the substrate) in our previous study.[1] Region iii is represented by a small cluster with a bit of a distorted helix and a loop that was found to be in the beta region but not forming a true strand.

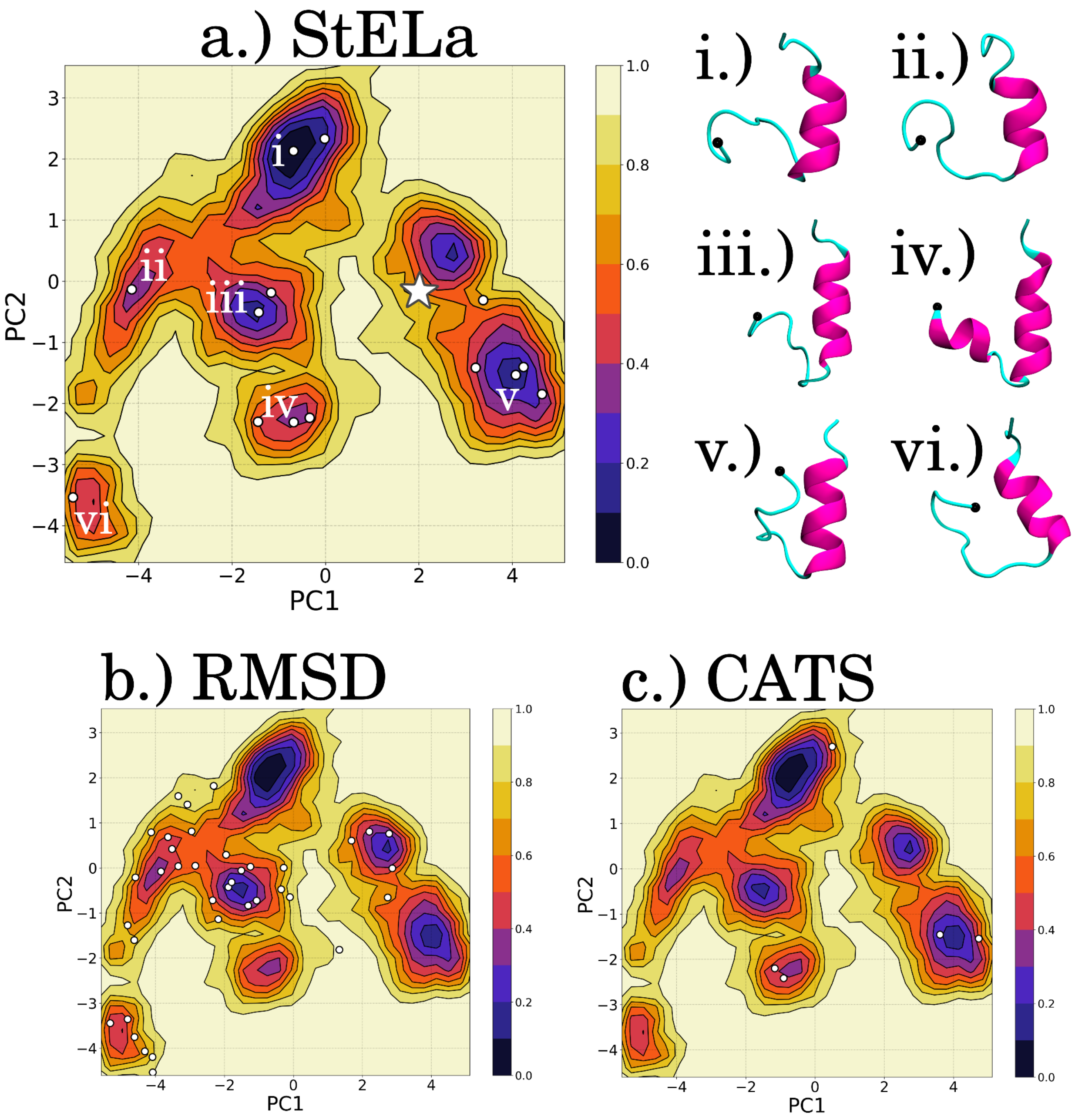


**Figure S15**: The representative structures, made with VMD,[6] for the identified clusters for the HBD tip of the **Monomer APO** set up are plotted on the FEL in PC1/PC2 space for each method: (a) StELa, (b) RMSD and (c) CATS+.[1,7] The white star in (a) indicates where the starting structure is found in the beginning of the simulation. Representative structures (i-vi) were extracted to represent the minima indicated in (a). The N-terminal end of the structure is indicated with a black bead. Neither StELa nor CATS+ identified a cluster in one of the minima between i and v so a representative for it is not provided.

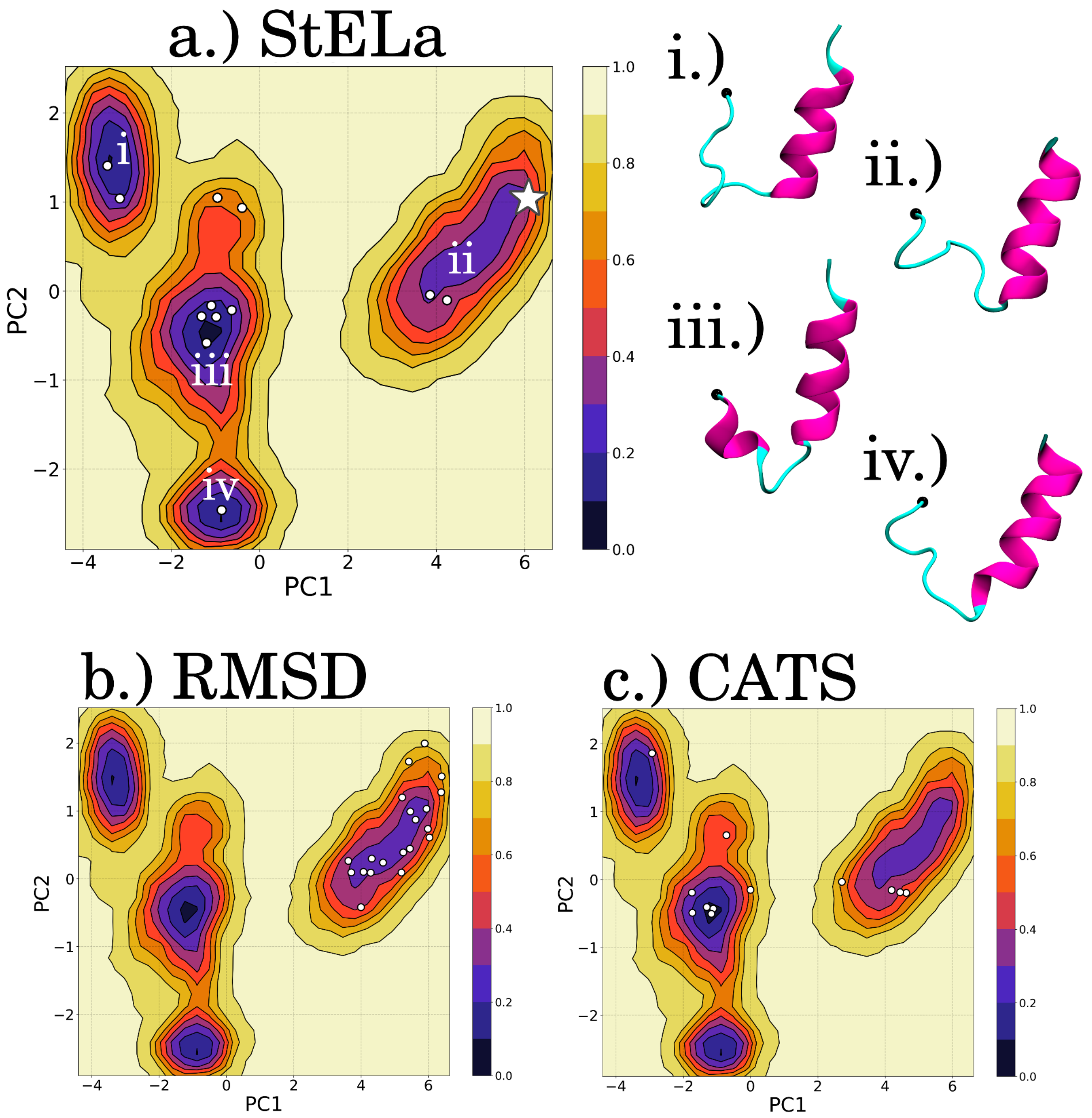


**Figure S16**: The representative structures, made with VMD,[6] for the identified clusters for the HBD tip of **Protomer A** from the **Ring-ABC-COMPLEX** set up are plotted on the FEL in PC1/PC2 space for each method: (a) StELa, (b) RMSD and (c) CATS+.[1,7] The white star in (a) indicates where the starting structure is found in the beginning of the simulation. Representative structures (i-vi) were extracted to represent the minima indicated in (a). The N-terminal end of the structure is indicated with a black bead.

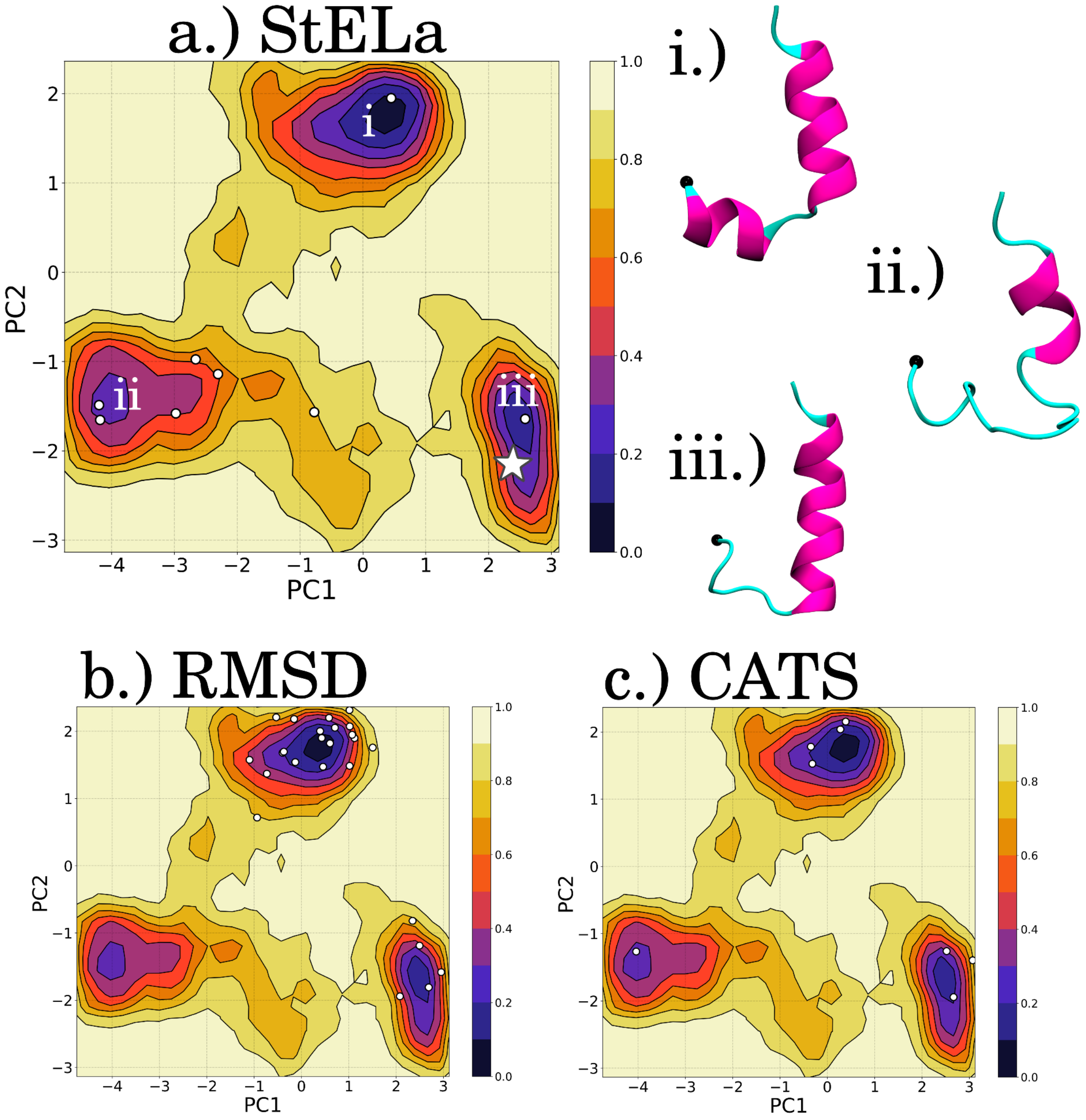


**Figure S17**: The representative structures, made with VMD,[6] for the identified clusters for the HBD tip of **Protomer A** from the **Ring-ABC-APO** set up are plotted on the FEL in PC1/PC2 space for each method: (a) StELa, (b) RMSD and (c) CATS+.[1,7] The white star in (a) indicates where the starting structure is found in the beginning of the simulation. Representative structures (i-iii) were extracted to represent the minima indicated in (a). The N-terminal end of the structure is indicated with a black bead.

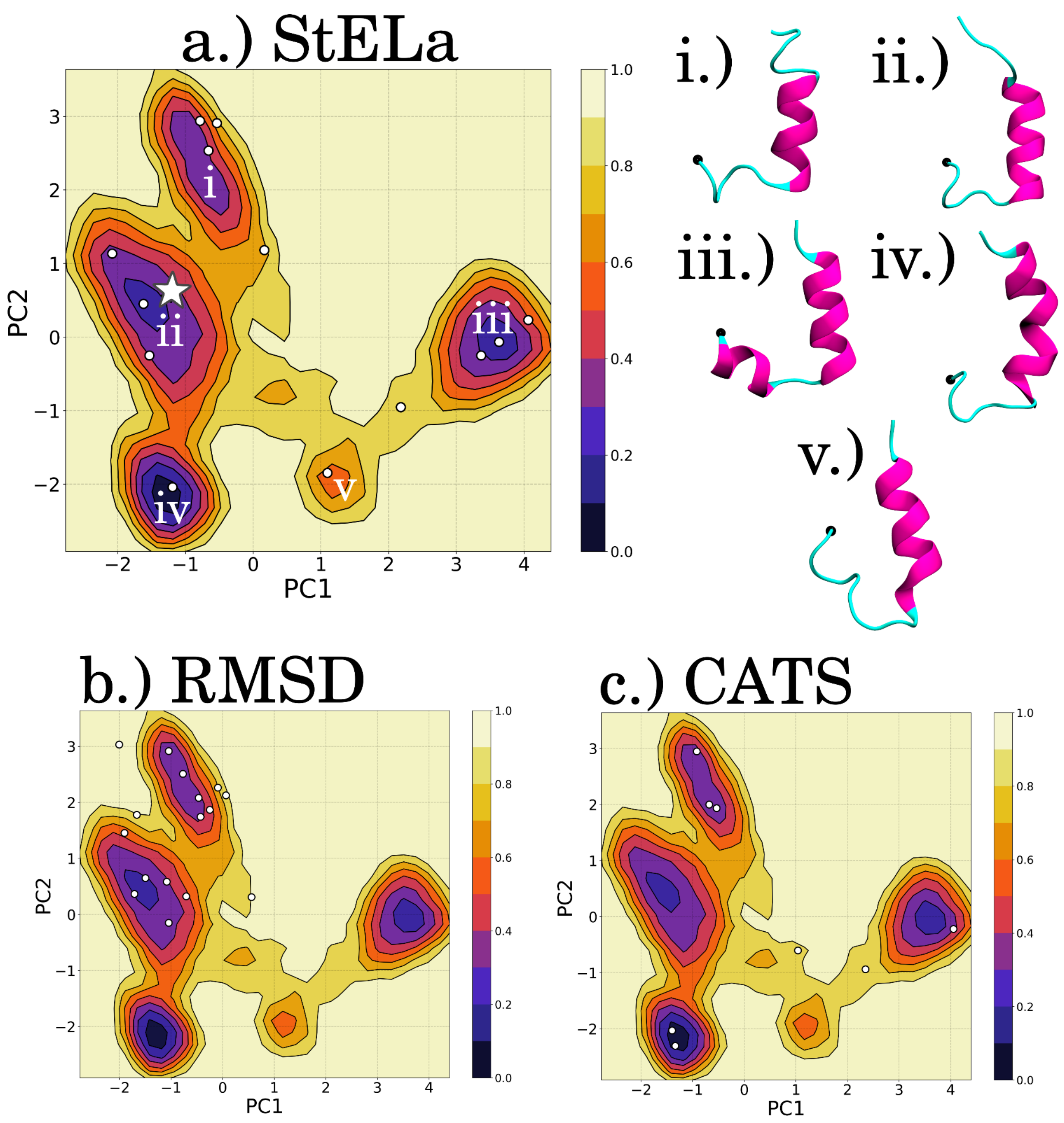


**Figure S18**: The representative structures, made with VMD,[6] for the identified clusters for the HBD tip of **Protomer B** from the **Ring-ABC-COMPLEX** set up are plotted on the FEL in PC1/PC2 space for each method: (a) StELa, (b) RMSD and (c) CATS+.[1,7] The white star in (a) indicates where the starting structure is found in the beginning of the simulation. Representative structures (i-v) were extracted to represent the minima indicated in (a). The N-terminal end of the structure is indicated with a black bead.

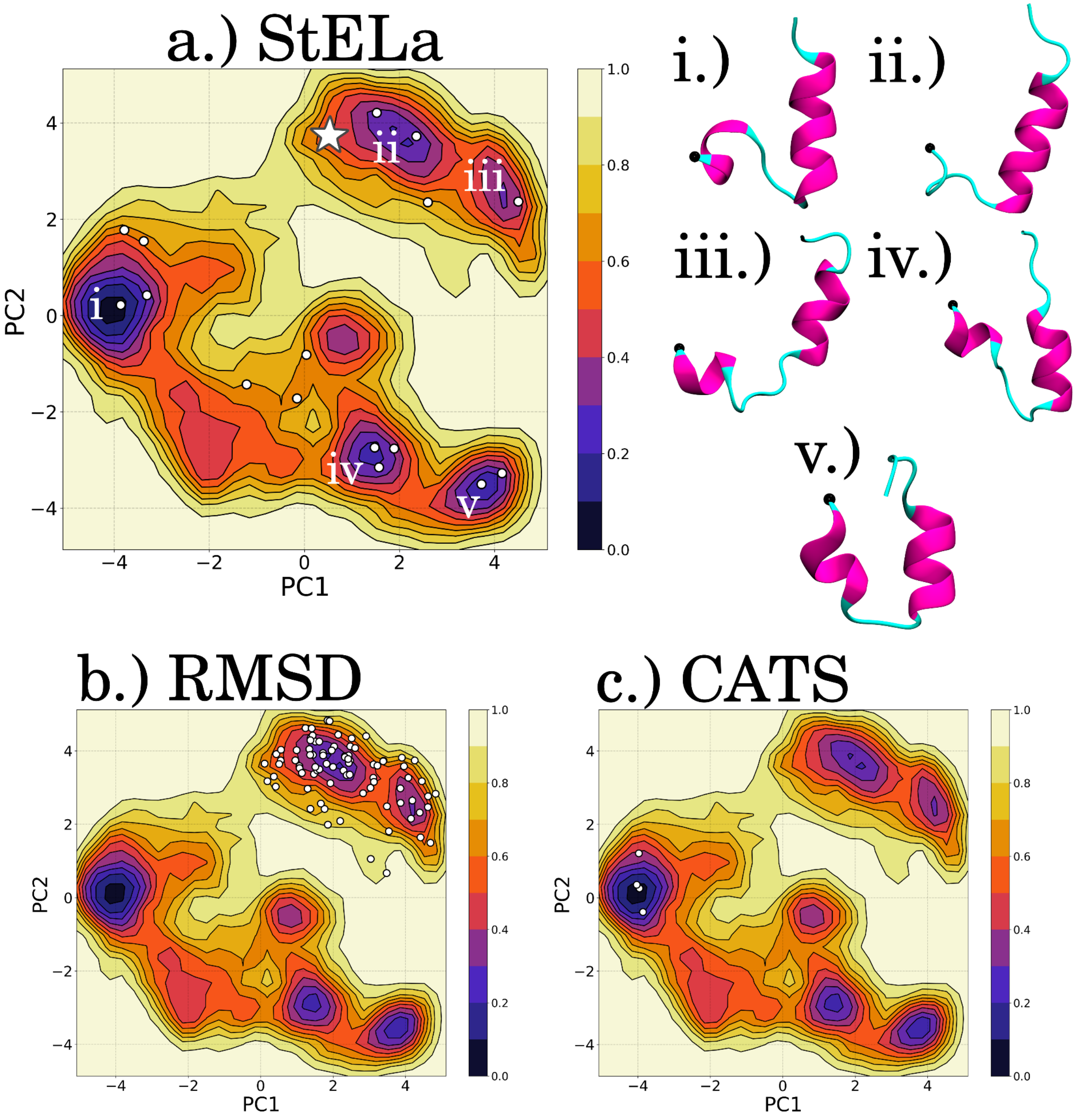


**Figure S19**: The representative structures, made with VMD,[6] for the identified clusters for the HBD tip of **Protomer C** from the **Ring-ABC-COMPLEX** set up are plotted on the FEL in PC1/PC2 space for each method: (a) StELa, (b) RMSD and (c) CATS+.[1,7] The white star in (a) indicates where the starting structure is found in the beginning of the simulation. Representative structures (i-v) were extracted to represent the minima indicated in (a). The N-terminal end of the structure is indicated with a black bead. There is a more shallow region that is not isolated by any of the methods and therefore is not represented.

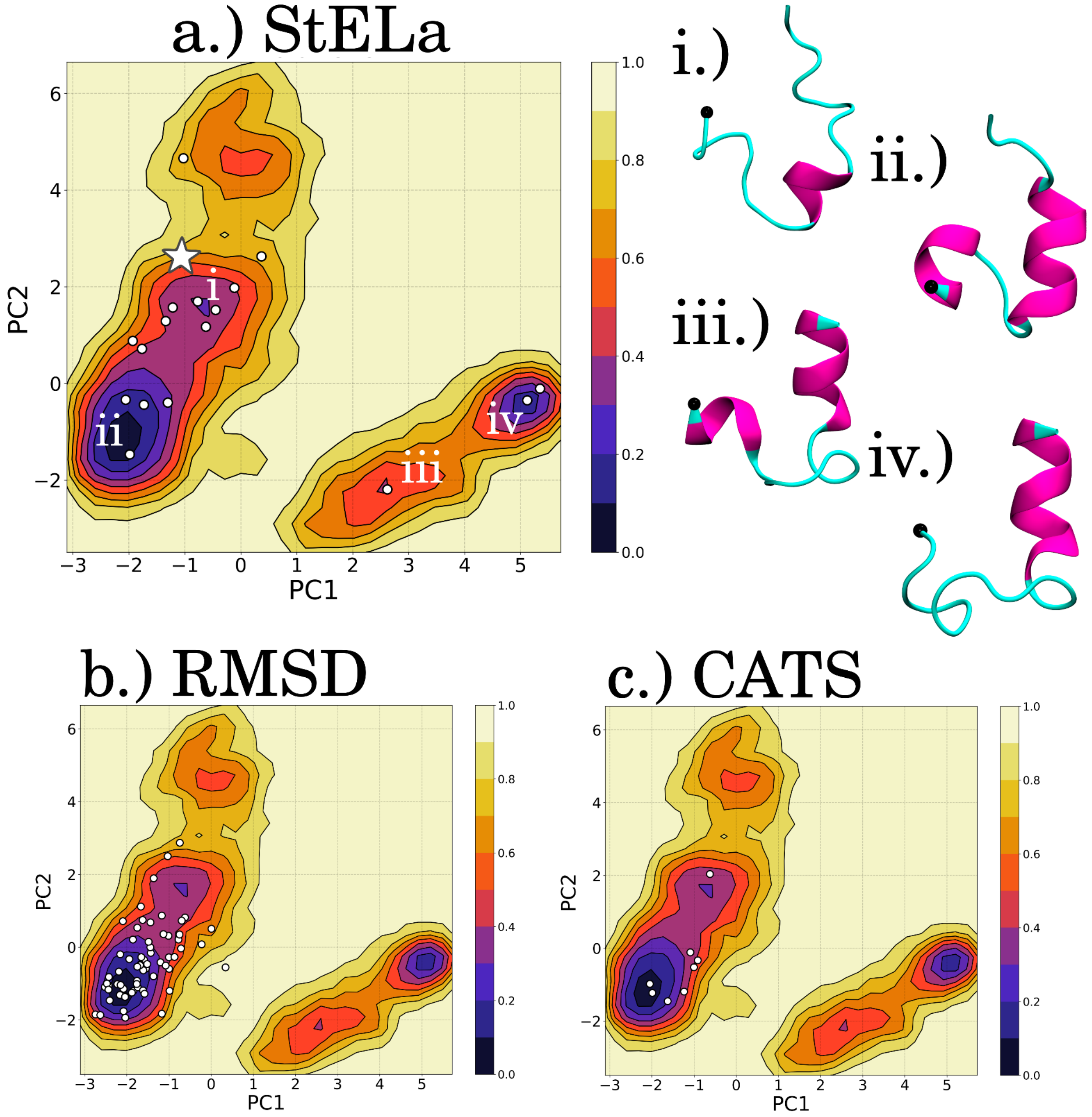


**Figure S20**: The representative structures, made with VMD,[6] for the identified clusters for the HBD tip of **Protomer C** from the **Ring-ABC-APO** set up are plotted on the FEL in PC1/PC2 space for each method: (a) StELa, (b) RMSD and (c) CATS+.[1,7] The white star in (a) indicates where the starting structure is found in the beginning of the simulation. Representative structures (i-iv) were extracted to represent the minima indicated in (a). The N-terminal end of the structure is indicated with a black bead. There is a more shallow region that is not isolated by any of the methods and therefore is not represented.

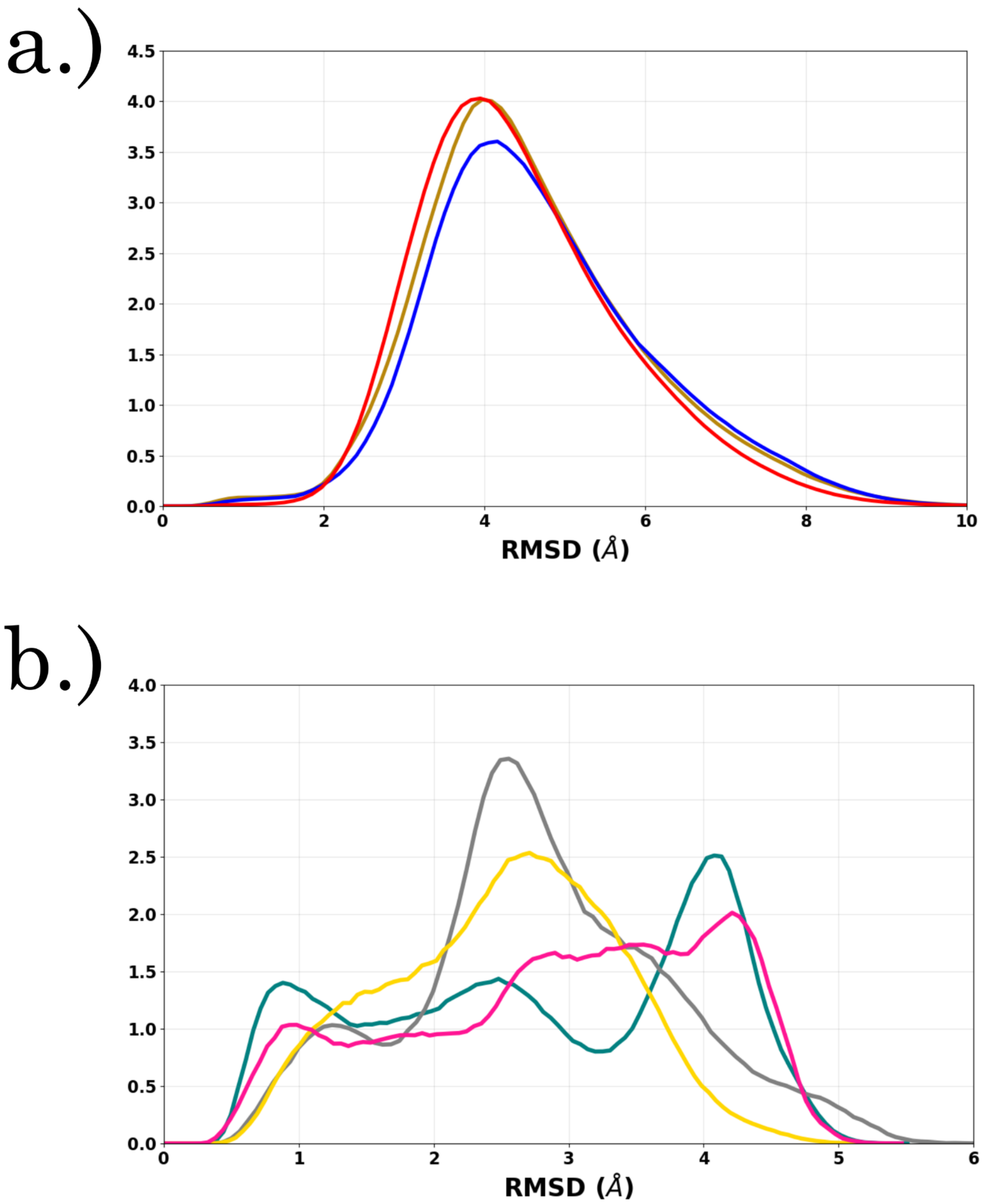


**Figure S21:** (a) Depicts the RMSD distributions extracted with the RMSD-based algorithm[10] for the R2 fragment of Tau in each of the conditions included in this study (**TMAO** - yellow, **Water** - blue, **Urea** - red). From this plot, we identify that there is not a clear separation of the distributions indicating RMSD may not be a descriptive collective variable for clustering secondary structures. (b) Shows the RMSD distributions extracted with the RMSD-based algorithm for the HBD tip of the Katanin Monomers in each of the ligand states included in this study (**COMPLEX** - teal, **NUC** - yellow, **SUB** - pink, **APO** - gray).

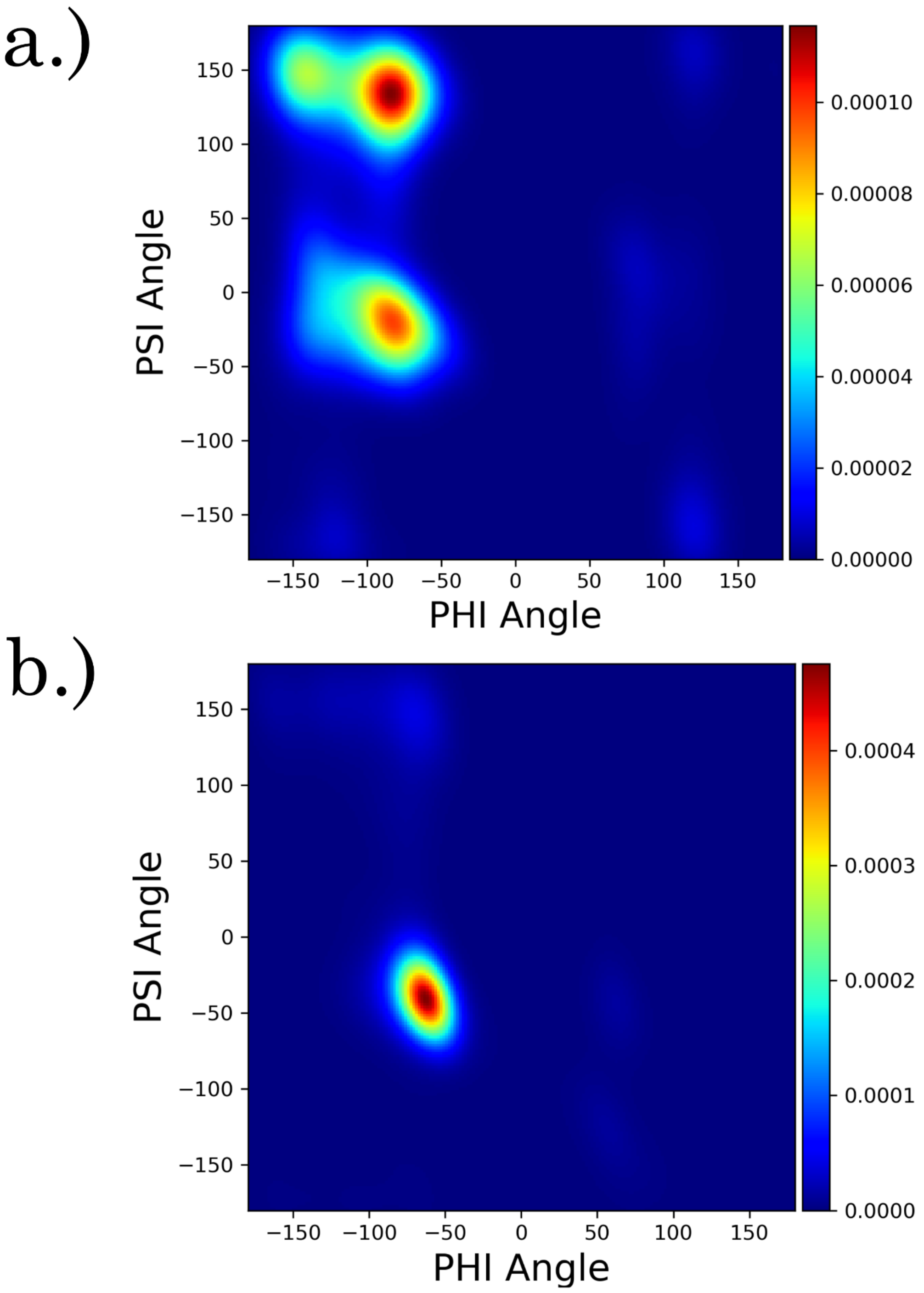


**Figure S22**: The Ramachandran plot colored according to the KDE determined for (a) the R2/Tau fragment in **Water** and for (b) the HBD tip of the **monomer-COMPLEX** setup. The R2/Tau fragment populates more of the PHI/PSI space in general, in particular the region associated with beta strands and sheets in comparison to the HBD tip fragment that was found to be nearly exclusively helical.[11,12]
